# Zebrafish eye development and function is affected by crude oil exposure – line of evidence from underlying molecular effects to behavioral consequences

**DOI:** 10.1101/2024.01.23.576882

**Authors:** Leonie K. Mueller, Sarah Johann, Bernd Denecke, Ali T. Abdallah, Markus Hecker, Henner Hollert, Thomas-Benjamin Seiler

**Affiliations:** Department of Ecosystem Analysis, Institute of Environmental Research, ABBt – Aachen Biology and Biotechnology, RWTH Aachen University, Aachen, Germany; Altertox, Brussels, Belgium; Department of Evolutionary Ecology and Environmental Toxicology, Goethe University Frankfurt, Frankfurt am Main, Germany; IZKF Aachen, University Hospital, RWTH Aachen University, Aachen, Germany; Cluster of Excellence on Cellular Stress Responses in Aging-Associated Diseases (CECAD), University of Cologne, Joseph-Stelzmann-Str. 26, 50931 Cologne, Germany; School of the Environment & Sustainability, Toxicology Centre, University of Saskatchewan, Saskatoon, Canada; State Key Laboratory of Pollution Control and Resource Reuse, School of the Environment, Nanjing University, Nanjing, China; Fraunhofer Institute for Molecular Biology and Applied Ecology IME, Department Ecotoxicology across environmental media, Schmallenberg and Frankfurt; LOEWE Centre for Translational Biodiversity Genomics (LOEWE-TBG), Senckenberganlage 25, 60325 Frankfurt am Main, Germany; Hygiene-Institut des Ruhrgebiets, Gelsenkirchen, Germany

## Abstract

Crude oil can affect the normal eye development and swimming behavior of fish embryos. In this study 0-4 hours post fertilization (hpf) zebrafish (*Danio rerio*) embryos were exposed to low, sublethal concentrations of water-accommodated fractions (WAF) of a naphthenic North Sea crude oil alone and in combination with a third-generation chemical dispersant. Effects on eye development and function at the transcript, histological, morphological, and behavioral level were assessed relative to untreated control specimen at 119 hpf. Exposed embryos swam significantly less during the dark phases of the light/dark transition test. Eye size of zebrafish embryos was significantly smaller in groups exposed to crude oil WAFs. Zebrafish retina histology revealed that, among other structural alterations, the outer nuclear layer, containing photoreceptor cells relevant for normal visual functioning, was significantly thinner in exposed embryos. Transcriptome analysis revealed that several genes involved in eye development and function (opsins and crystallins) were dysregulated upon oil exposure. The molecular effects on gene expression and eye histology results form a novel line of evidence that explains the observed effects of crude oil on the swimming activity of exposed fish embryos. The present study is in line with recent findings highlighting the importance of ocular toxicity in developing embryos exposed to petroleum samples at environmentally relevant and non-lethal concentrations without visible malformations. The study suggests that oculotoxicity should be considered as a major toxicity pathway alongside to the well-established cardiotoxicity.

## 1. Introduction

Increasing global energy demands have resulted in the intensification of exploration and transportation of crude oil, which significantly increased the risk of oil spills over the past five decades. Between 1974 and 2010 approximately 1200 oil spills occurred (Eckle et al., 2012). Although the number of tanker accidents that released large amounts of oil into the environment has declined in the recent past (Chen et al., 2019; ITOPF, 2022), the risk of large oil spills remains, as demonstrated by several incidents from both oil exploration (e.g. Deepwater Horizon blowout 2010) and transportation activities (e.g. SANCHI tanker collision 2018, tanker spill at Qingdao port 2021). Once introduced into the environment the surface oil slick and the dispersion of oil droplets due to wind and waves releases many oil constituents dominated by lower molecular weight (LMW) polycyclic aromatic hydrocarbons (PAHs) into the water column (Gong et al., 2014) where they can affect aquatic organisms, including adult and early life stages of pelagic fishes. In addition to the potential acute adverse effects caused by these low molecular weight compounds, common crude oil constituents, including higher molecular weight PAHs, can persist in affected areas for decades and result in long-term effects in these ecosystems (Peterson et al., 2003; Reddy et al., 2002; Short et al., 2004). Early life stages of fish are particularly sensitive to the adverse effects of crude oil (Pasparakis et al., 2019; Xu et al., 2016) and the exposure is potentially high, as the toxic compounds accumulate in near surface waters where fish early life stages reside (Fodrie and Heck, 2011). Furthermore, the application of chemical dispersants commonly used in oil spill responses is a critical consideration for the toxicological assessment of such spills because they increase the dissolved and particulate oil fractions (Prince, 2015).

Cardiotoxicity has been described as the primary phenotype of crude oil toxicity in developing fish embryos (Incardona, 2017), which can be triggered via AhR-dependent and -independent molecular mechanisms. However, based on a review of the literature describing the adverse effects of the Deepwater Horizon oil on teleost fishes it has been concluded that oil toxicity in developing fish is a multi-organ effect (Pasparakis et al., 2019). In fact, a variety of effects of crude oil other than cardiotoxicity have been described in fish, including altered development of craniofacial and spinal structures (de Soysa et al., 2012; Corinne E. Hicken et al., 2011; Incardona et al., 2004; Khursigara et al., 2017), oxidative stress, disruption of energy metabolism and narcosis (Crowe et al., 2014; Gagnon and Holdway, 1999; Meador and Nahrgang, 2019; Nahrgang et al., 2010), which might not all be secondary effects resulting from cardiotoxicity. In particular, the impact on sensory organ development receives increasing attention in oil toxicity assessment. Pasparakis et al. (2019) highlighted the need for additional studies investigating the effects of oil constituents on the visual system in fish. Several recent studies focused on effects of individual AhR-agonists and crude oils on the development and normal function of the visual system in early life stages of different fish species (Li et al., 2020; Lie et al., 2019; Magnuson et al., 2020; Philibert et al., 2021; Xu et al., 2016). These studies found effects on the visual system at different levels of biological organization, e.g., the transcriptomic dysregulation of certain biological pathways relevant for eye development, eye morphology and structure or altered fish swimming behavior. However, attempts to assess the effects of crude oil exposure to fish across biological levels of organization trying to link molecular response patterns to behavioral alterations has been limited to a few studies. The underlying molecular mechanism linking crude oil or oil constituents (e.g. PAHs) to alterations of the visual system has been suggested to be attributed to AhR-responsive genes in the zebrafish (*Danio rerio*; Aluru et al., 2014). However, the exact molecular mechanisms are not well understood to date.

Hence, this study used whole transcriptome sequencing in combination with histological and swimming behavior investigations in early life stages of zebrafish to assess the underlying molecular mechanisms of naphthenic North Sea crude oil alone and in combination with a third-generation chemical dispersant. Within this context, unbiased total RNA sequencing was used as it been identified as a suitable tool in ecotoxicological risk assessment (Mehinto et al., 2012) allowing a connection to higher biological functions. As a behavioral endpoint the light/dark transition test was utilized that allows to measure the ability of the fish embryos to maintain normal reactions to a negative visual stimulus. Zebrafish larvae eye histology was selected to provide more insight into potential morphological alterations and to bridge the gap between the molecular and behavioral effects on the visual system.

## 2. Materials and Methods

### 2.1. Animals

Wild-type zebrafish from the West Aquarium strain (Bad Lauterburg, Germany) were used in this study. Breeding groups of 100 to 150 adult zebrafish ranging between 1 and 2 years of age were kept in recirculating 170 L tanks under flow-through conditions with a water exchange rate of 40 % per week equipped with biol-filters and UV sterilization. Fish were fed twice a day with dry flakes (TetraMin, Tetra GmbH, Melle, Germany) and larvae of *Artemia* spec. A constant day-night rhythm (14:10) and temperature (26 ±1 °C) was maintained. Spawning took place 30 min after the onset of light.

### 2.2. Preparation of water-accommodated fractions

A naphthenic North Sea crude oil and the commercially available third generation chemical dispersant Finasol OSR 51 (Total, Paris la Défense, France) were used. The low energy water accommodated fractions (LEWAFs) and chemically enhanced water accommodated fractions (CEWAFs) were prepared according to Singer et al. (2000) with minor modifications (Johann et al. 2020). Briefly, LEWAFs and CEWAFs were prepared in aspirator glass flasks (500 mL) by application of oil or a 1:10 (w/w) dispersant-oil mixture on the surface of formulated fish water (294.0 mg/L CaCl_2_ · 2H_2_O; 5.5 mg/L KCl; 123.3 mg/L, MgSO4 · 7H_2_O; 63.0 mg/L NaHCO_3_, pH 7-8) at an oil-to-water ratio of 1:50 (w/v, LEWAF) and 1:200 (w/v, CEWAF), respectively. Oil-to-water ratios were based on EU project GRACE agreement and kept constant for every study performed within this context (Jørgensen et al., 2019). The LEWAF setup was stirred with low energy avoiding a vortex in the water phase while the CEWAF was stirred at higher speeds to create a 25 % vortex of the water phase. LEWAFs and CEWAFs were stirred for 40 h at 10 °C, as specified by the EU project GRACE (Jørgensen et al., 2019). After 1 h settling time, LEWAFs and CEWAFs were then drained and diluted in formulated fish water. Air-sealed 10 mL glass vials with low head space were used as exposure vessels. LEWAF and CEWAF exposure solutions were warmed up to 26 °C before usage. The exposure concentrations prepared were 12.5 % (1:400 dilution, LEWAF) and 0.8 % (1:25000, CEWAF) of 100% stock solutions, respectively. The exposure solutions were selected based on 10 % effect concentrations (EC10) derived from preceding fish acute embryo toxicity tests according to OECD guideline 236 with identical oil batches and experimental setups (Johann et al., 2020).

### 2.3. Exposure regime

The exposure of zebrafish embryos was performed according to OECD guideline 236 (OECD 2013) with minor modifications. Briefly, for each treatment 50 embryos were used for transcriptomics analyses, 40 for qPCR verification, 10 embryos for histological analyses, and 30 embryos for the behavioral tests. Embryos were transferred to exposure solutions shortly after fertilization (10 embryos per 10 mL glass vial). Embryos were incubated at 26 °C using a semi-static approach with periodic medium exchange (every 24 h) as described in detail in Johann et al. (2020). Formulated fish water was prepared, aerated, and warmed up one day before using. The pH of all media was adjusted to values between 7.0 and 8.0 (using 0.1 M HCl or 0.1 M NaOH). For all exposure regimes 4 independent experiments were performed.

### 2.4. Chemical analysis

Exposure concentrations of 18 priority US EPA PAHs (for a complete list of target compounds see SI 1 Table SI 1) were measured daily according to the methods detailed in (Potter and Pawliszyn, 1994) in samples of the freshly prepared LEWAFs and in samples from the LEWAFs that were removed during the daily medium exchange. The chemical analysis was included to assess the stability of exposure concentrations of polar organic compounds in the test vessels. Target PAHs were extracted directly from the medium using 30 µm silicone coated solid phase micro extraction (SPME) fibers (Supelco, Sigma-Aldrich Corp.). The concentrations of selected PAHs in the CEWAF were not measured since the presence of the dispersant and oil droplets would have interacted with the loading rates and potentially destroyed the silicone coated fibers. Perdeuterated internal standard PAHs were added to the samples prior to extraction. External (S-4008-100-T) and perdeuterated internal standards (S-4124-200-T) were purchased from Chiron (Chiron AS, Trondheim, Norway). Extraction was conducted for 2 h to enable quantification of low concentrations of target PAHs in the test medium. Loaded SPME fibers were removed from the test medium, carefully dried using paper tissues and immediately analyzed using an Agilent Technologies GC system (7890 A GC System and 5975 C inert XL MSD with Triple-Axis-Detector, Agilent Technologies Deutschland GmbH) (Potter et al. 1994).

### 2.5. Exposure termination

Exposure of embryos was terminated at 119 hpf, staying within non-animal testing according to European legislation (EU, 2010, 1986; Strähle et al., 2012). All embryos were anesthetized using saturated benzocaine solution and transferred to 1.5 ml micro test tubes (VWR International). Embryos for RNA sequencing were washed twice with cold phosphate buffered saline (PBS, Sigma-Aldrich Corp.) and excess solution was replaced by 200 µl of RNAlater (Sigma-Aldrich Corp.). Subsequently, embryos were snap-frozen in liquid nitrogen.

### 2.6. Behavioral assay

For the assessment of differences in behavioural responses, 30 embryos from each treatment were collect in the morning of the day of termination of the exposure for transcriptome analysis at 119 hpf. For the behavioral assays, embryos were individually transferred to a 96 well plate (STARLAB GmbH) and subsequently kept in the incubator for 30 min to allow for recovery from the handling stress prior to initiation of the test. The behavioral test was conducted using a DanioVision observation chamber and EthoVision tracking software (Noldus Information Technology). Embryo swimming activity was assessed by a light dark transition test and tracked for 50 min. After an initial acclimatization period of 10 min in the dark, transition between light and dark phases was assessed (10:10 min). After completion of the behavior test, embryos were euthanized by prolonged immersion in a benzocaine-ethanol solution. Differences in embryo moving patterns were analyzed in 1-min duration time blocks.

### 2.7. Body length measurements and eye histology

At the end of the exposure experiment zebrafish embryos were euthanized using saturated benzocaine solution in ultra-pure water (< 1 g/L) and rinsed twice with PBS. Afterwards, embryos were transferred to 1.5 mL tubes and fixed in 4 % paraformaldehyde solution (PFA) over night at 4 °C followed by a serial dehydration in methanol (50 - 100 % MeoH). Embryos were stored at -20 °C in 100 % MeOH until further usage.

For the body measurements 10 fixed embryos for each of the 4 replicates per treatment (n=40 per treatment) were orientated in lateral plane and inspected using an inverted microscope (Nikon Eclipse Ti2, Nikon GmbH, Düsseldorf, Germany) with 2x magnification. Generated images of embryos were analyzed for total body length and eye size of the embryos with the associated software NIS Elements BR (V.5.11, Nikon GmbH) as described before (Baumann et al., 2016). The relative eye size of 10 embryos per treatment was calculated as the ratio of eye diameter over total body length.

For histological analysis of zebrafish eyes 8 MEOH-fixed embryos per treatment were embedded and orientated in agarose molds (2 % agarose). After drying, the agarose molds were transferred to histology cassettes (Simport Scientific, VWR International), and processed in a tissue processor (MTM, Slee medical GmbH) using an over-night program with a series of ethanol (70 - 100 %) and xylol dehydration followed by paraffin embedding. Coronal sections (3 µm) of zebrafish embryo heads were produced with a microtome (pfm slide 4003 E, Pfm Medical GmbH) using the following procedure: sections of 10 µm thickness were produced until the lens was visible and discarded. Afterwards, 10 serial sections of 3 µm thickness per embryo were prepared to guarantee the inclusion of slides with visible optic nerve. Slides were dried at 37 °C over-night and stained with hematoxylin/eosin (H/E) using an automated slide stainer (Gemini AS, Thermofischer Scientific).

An inverse microscope (Nikon Eclipse Ti2) with 20x magnification and the associated software NIS Elements BR (V.5.11, Nikon GmbH) were used to analyze the H/E-stained slides of 8 embryos per treatment (2 embryos per biological replicate). Only coronal sections showing the plane with optic nerve were used for measurements to guarantee comparable section depth. The diameter of the 5 retinal cell layers at 8 different locations as well as the lens diameter were measured for both eyes of individual embryos, if possible. Otherwise, one eye per embryo was evaluated. Furthermore, the pigmentation of the retinal pigment epithelium was quantified as described in Baumann et al. (2016). Images were converted to an 8-bit grey scale from 0 (black) to 250 (white). The intensity of the pigment epithelium was measured as the mean intensity of drawn areas (2 -10 µm^2^) at the 8 locations for retinal layer measurement.

### 2.8. RNA extraction, cDNA synthesis and Illumina sequencing

Total RNA from pools of 50 zebrafish embryos (119 hpf) for each of 4 independent biological replicate was isolated using the Maxwell RSC miRNA tissue kit according to the manufacturer’s instructions. RNA samples were assessed for quality (integrity) with the TapeStation 4200 using the Agilent RNA ScreenTape Assay kit (all eRINs were >9). Quantification was performed using the Quantus Fluorometer and QuantiFluor RNA Dye (Promega GmbH, Mannheim, Germany). All cDNA libraries were generated from 1 µg total RNA using the TrueSeq Stranded Total RNA Library Preparation Kit with the Ribo-Zero Gold Kit according to the manufacturer’
ss instructions (Illumina Inc., San Diego, CA, USA). Quality and quantity of the cDNA libraries were assessed using a TapeStation 4200 (D1000 screen tape assay) and a Quantus Fluorometer, respectively. The libraries were run on an Illumina NextSeq 500 platform using the High Output 150 cycles Kit (2 x 75 bp paired end reads) resulting in a sequencing depth between 46.3 and 57 million reads per sample.

### 2.9. Validation by RT-qPCR

To quantitatively confirm the results of the Illumina sequencing real time quantitative PCR (RT-qPCR) was performed with a set of genes that were significantly dysregulated. The primers for the selected transcripts (SI 1 Table SI 2) were designed using the primer designing tool of the National Institutes of Health, verified via oligo DNA product sequencing, and ordered from Eurofins Genomics (Ebersberg, Germany). Total RNA concentrations were determined at 260 nm using a NanoDrop ND-1000 134 Spectro-photometer (Thermo Fisher Scientific Inc.). Total cDNA for RT-qPCR was generated from 1.5 µg total RNA using a random primer mix and M-MuLV reverse transcriptase (New England Biolabs) according to the manufacturer’s protocol. RT-qPCR was performed in 96-well plates (PCR-96-LP-AB-C, Axygen Inc.) with SYBR^®^ green Real-Time PCR Master mix (Thermo Fisher Scientific Inc.). The PCR program included an enzyme activation step at 95 °C (10 min) and 40 cycles of 95°C (15 sec) and 60 - 62°C depending on the target gene (SI 1 Table SI 2) (1 min). Additionally, a melting curve was added to ensure target specificity and single peak amplification. Target gene expression levels were normalized to the expression of the reference gene elongation factor 1 (eef1a1). The mathematical model for relative quantification by Pfaffl et al. (2001) was used 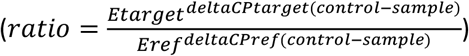

### 2.10. Statistical analysis

#### Statistical analysis of swimming behavior and size measurements

For statistical analysis of the data from the behavioral assays, the results of minute 25, representing peak activity levels during the dark phase, were compared between control and treatment groups. Results for the overall swimming distance were tested for normal distribution of the data using the Shapiro-Wilk test. Since the data was not normally distributed, Kruskall Wallis test with subsequent pairwise Wilcoxon rank sum test was used to test for significant differences between control and treatment groups (p<0.05).

To identify significant differences in whole embryo size and retina measurements of exposed embryos compared to control, statistical analysis using One Way ANOVA with Dunnett’s post hoc test after verifying normal distribution (Shapiro-Wilk test) and equal variance (Lavene test) was performed (p<0.05). For data not normal distributed or not equal in variance non-parametric Kruskal Wallis test with Dunn’s post-hoc test was performed (p<0.05).

#### Functional annotation and statistical analysis of transcriptome results

RNA-Seq data was analyzed using an in-house pipeline embedded in the QuickNGS workflow management system (Wagle et al., 2015). Generation of the fastq files and adapter removal were completed using the Illumina software bcl2fastq (https://support.illumina.com/sequencing/sequencing_software/bcl2fastq-conversion-software.html). Quality control checks of the RNA-Seq data were done with fastqc available online (http://www.bioinformatics.babraham.ac.uk/projects/fastqc). Subsequently, the reads were aligned to the mm10 reference genome sequence using STAR 2.5.2b (Dobin et al., 2013) with default parameters. In the quantification step, reads were counted with featureCounts (Liao et al., 2014) from the subread package v1.5.1 available online (http://subread.sourceforge.net). Afterwards, differential expression analysis was performed in R using the DESeq2 package (Love et al., 2014) selecting the local regression parameter and otherwise default parameters.

Functional annotation analysis was performed by GO- and KEGG-Pathway-enrichment analyses for each of the test conditions (LEWAF and CEWAF). The analysis was performed using Cytoscape (Shannon et al., 2003) and the plug-in ClueGO (Bindea et al., 2009) against the Gene Ontology and KEGG databases. Functional annotation of significantly differentially expressed genes with a 1.5 fold-change cut off was performed by uploading gene identifiers of differentially expressed genes to ClueGO. Genes were functionally annotated by assigning GO terms for the domains biological process, molecular function, and cellular component and by assigning genes to KEGG Pathways. The annotation network was created using the default settings of ClueGO. To ensure an up-to-date functional analysis, latest ontology and annotation sources were downloaded on 16.06.2023. All terms with a p-value <0.05 were considered to be significantly regulated.

## 3. Results & Discussion

### 3.1. PAH exposure regime

Out of the 18 target PAHs analyzed in the exposure medium, 3 PAHs were detected and quantified in the LEWAF samples collected before and after medium changes (SI 1 Table SI 3). Naphthalene (2-ring) was the most dominant of the analyzed PAHs in the LEWAF and was detected at concentrations between 17.9 and 33.2 μg/L in fresh exposure medium and between 17.1 and 25.0 μg/L in 1-day old exposure medium. Fluorene (3-ring) was detected at concentrations between 0.39 and 0.46 μg/L in fresh exposure medium and between 0.28 and 0.33 μg/L in 1-day old exposure medium. Finally, phenanthrene (3-ring) was detected at concentrations between 0.30 and 0.40 μg/L in fresh exposure medium and between 0.16 and 0.25 μg/L in 1-day old exposure medium. PAHs of higher molecular weights were either below the detection limit, or not present in the LEWAF samples. While several studies have shown that 3-ring PAHs in particular were able to cause adverse effects in fish embryos (Incardona, 2017), alkylated and heterocyclic derivates as well as other crude oil constituents not analyzed within the current setup could also have led to the observed effects discussed below (Bornstein et al., 2014; Martin et al., 2014).

The results showed that conditions during the experiment were relatively constant and that only little loss of freely available compounds occurred over the course of 24 h (SI 1 Table SI 3). PAH concentrations in the present exposure solutions (∑PAHs 18 – 34 µg/L) were in upper concentration ranges found in the water column after large oil spills (Echols et al., 2015; Tronczyński et al., 2004). The composition pattern with dominant low molecular weight and no to low concentrations of high molecular weight PAHs reflects a realistic scenario, since naphthalenes, phenanthrenes and alkyl-homologues have been identified most dominant dissolved PAHs in post-spill water column (Tronczyński et al., 2004). Hence, the exposure to the relatively high but still environmentally relevant exposure concentrations represents a realistic worst-case scenario.

### 3.2. Morphological and behavioral alterations

#### Eye histology

Exposure to the crude and chemically dispersed WAFs led to significant morphological changes of the eyes in developing zebrafish embryos. Both LEWAF and CEWAF treatments resulted in significantly reduced eye diameter compared to the control while total body length was not altered (Figure 1). Normalized eye size in WAF exposed embryos confirmed that the treatment led to significantly smaller eyes and was not due to overall smaller individuals caused by, e.g., developmental delay or general growth reduction.

**Figure 1.**
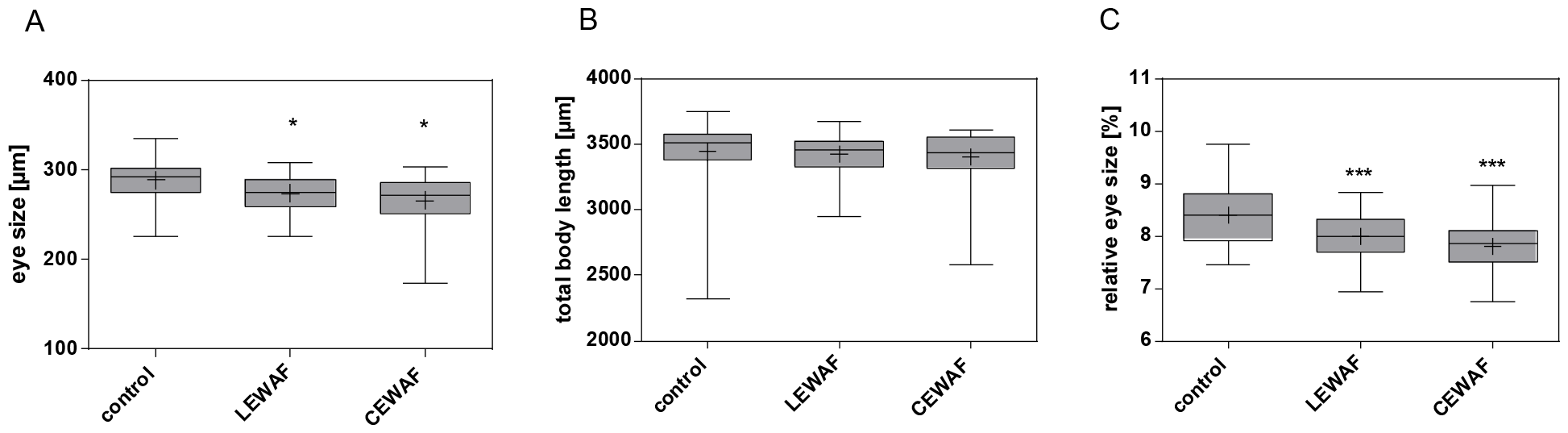
Whole larvae eye and body length of 119 hpf zebrafish embryos exposed to WAF dilutions of a naphthenic North Sea crude oil and the dispersant Finasol OSR 51. Zebrafish embryos were chronically exposed to WAF dilutions of crude oil only (LEWAF) and dispersed crude oil (CEWAF) at sublethal effect concentrations (EC10). Data are represented as mean (+) and median (line) with boxes showing the 25 – 75 percentile and whiskers the min to max values, respectively (n=40 per treatment). Relative eye size was calculated as the ratio of eye diameter over total body length. One Way ANOVA with Dunnett’s post hoc test was used for statistical analysis. In case normal distribution or equal variance tests failed, non-parametric Kruskal-Wallis One Way ANOVA on ranks with Dunn’s post hoc test was used. Asterisk indicate statistically significance of exposure groups compared to control group (*p<0.05, ** p<0.01, ***p<0.001).

Overall, reduced eye size in fish embryos exposed to oil has been reported for different fish species (Lie et al., 2019; Magnuson et al., 2020). In a similar study, effects on the eye diameter and body length were found in 96 hpf zebrafish embryos that were exposed to different crude oil WAF dilutions (Magnuson et al., 2020). However, in contrast to the present study, they found no significant effects on the relative eye size to body length. Though the total PAH concentration was in the range of the present study (13-90 µg/L) potential underlying reasons for the differences between both studies could be related to the oil types and corresponding composition with more 4-ring PAHs, and it is well known that every oil type can have unique adverse effects (e.g. Philibert et al., 2021).

The reduced eye sizes in our study were further attributed to the reduced diameter of particular cellular layers of the zebrafish retina investigated in H/E-stained coronal sections (Figure 2; details different layers see SI 1 Figure SI 1).

**Figure 2.**
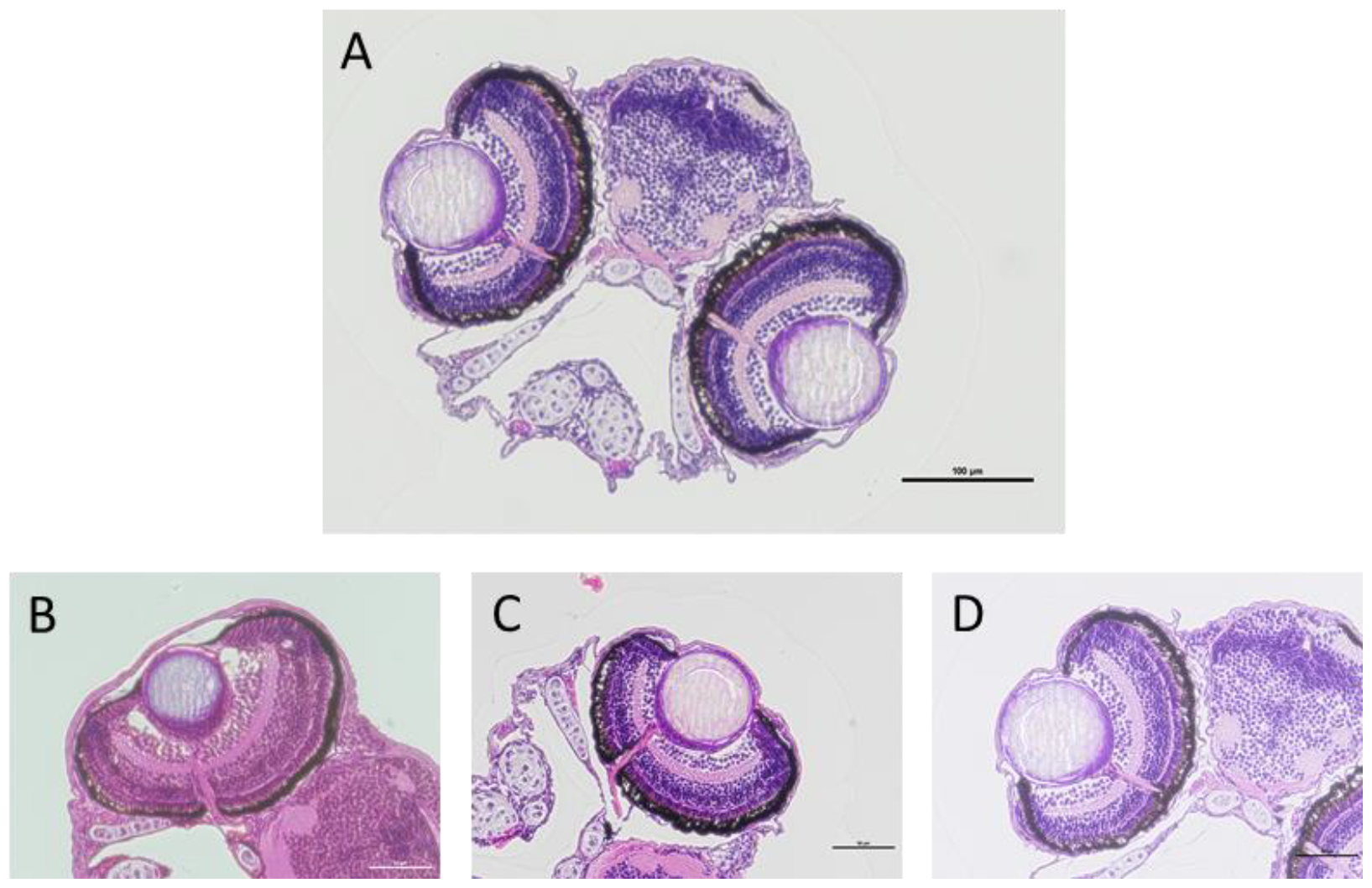
Coronal sections of 119 hpf zebrafish embryos exposed to WAF dilutions of crude oil only (LEWAF) and dispersed crude oil (CEWAF) at sublethal effect concentrations (EC10). H/E staining. A) 10x magnification of CEWAF treated embryos B – D) 20x magnification of control, LEWAF and CEWAF treatment. Scale bars= 100 µm (A), 50 µm (B-D).

The outer nuclear layer and the ganglion cell layer of the retina were significantly thinner in both treatment groups compared to the untreated control, while the lens diameter was significantly thinner in embryos exposed to chemically dispersed crude oil only (Figure 3). The plexiform layer was not found to be altered in WAF exposed embryos, while the inner nuclear layer was significantly thinner in LEWAF exposed embryos only. The outer nuclear layer contains the photoreceptor cells, thus, the reduced size of this specific compartment of the retina suggests structural alterations of the inner and outer segments of the photoreceptors in the eyes of WAF treated embryos.

**Figure 3.**
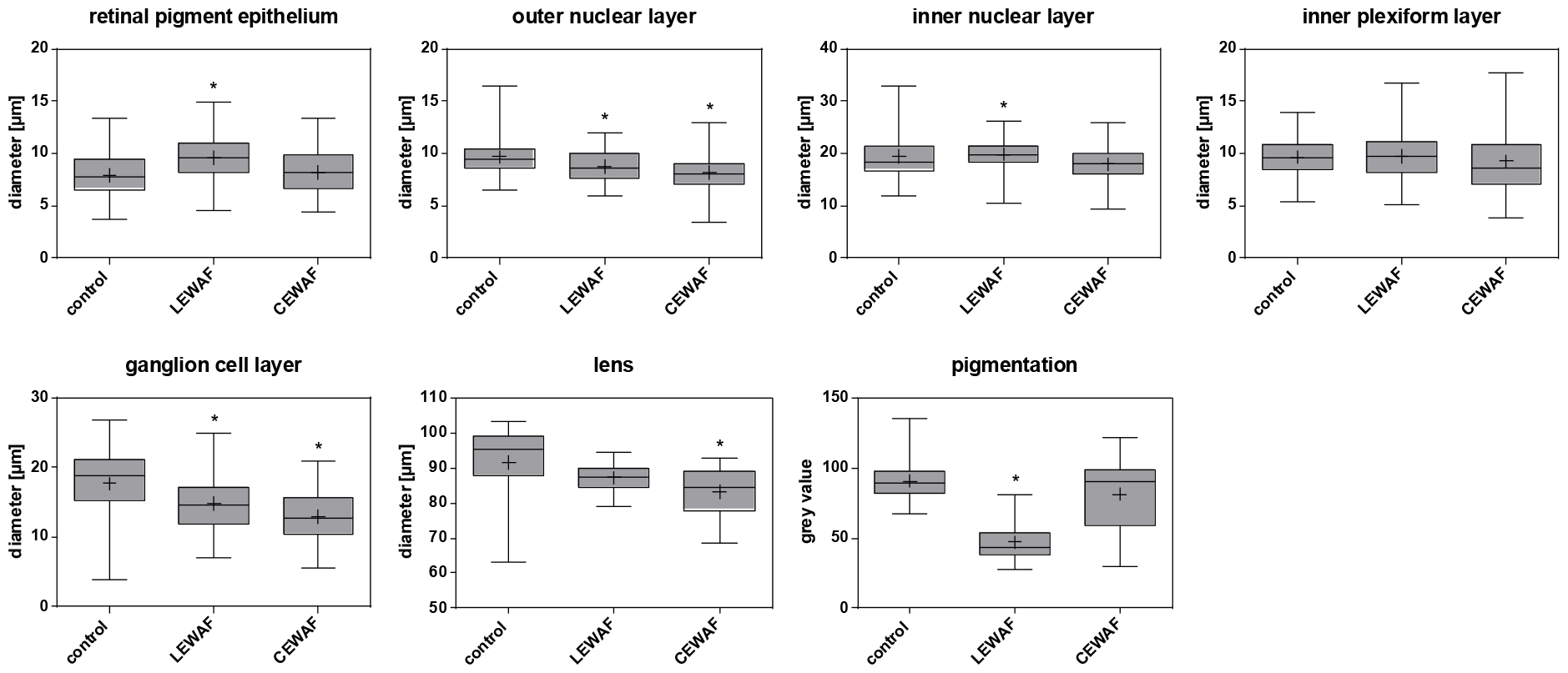
Changes in retinal layers and pigmentation of 119 hpf zebrafish embryos exposed to WAF dilutions of a naphthenic North Sea crude oil and the dispersant Finasol OSR51. Zebrafish embryos were chronically exposed to WAF dilutions of crude oil only (LEWAF) and dispersed crude oil (CEWAF) at sublethal effect concentrations (EC10). Data are represented as mean (+) and median (line) with boxes showing the 25 – 75 percentile and whiskers the min to max values, respectively (n=9-11 per treatment). As data were not normal distributed, non-parametric Kruskal-Wallis One Way ANOVA on ranks with Dunn’s post hoc test was performed to identify significant differences compared to control (p<0.05) indicated by Asterisks.

The photoreceptor cells of the zebrafish retina are organized in regular patterns (photoreceptor mosaic) (Fadool and Dowling, 2008). From H/E-stained histological sections of the samples in the present study no alterations in the mosaic were observed in crude oil exposed embryos. Gaps or random clustering would have resulted in under-representation or over sampling of information in those regions of the visual field (Fadool and Dowling, 2008). However, the structural information of simple H/E staining is limited. Immune labelling of specific cell types in the retina would provide more details. Furthermore, it has to be considered that the zebrafish retina is still developing in 119 hpf zebrafish embryos, and hence, the mosaic patterns are less pronounced compared to adults (Fadool and Dowling, 2008).

Interestingly, the retinal pigment epithelium (RPE) of LEWAF exposed zebrafish embryos was significantly thicker and stronger pigmented compared to control embryos, while CEWAF exposure did not lead to significant effects on pigmentation. This observation is in contrast to expectations, since previous studies investigating the impact of chemical exposure or nutrient alterations on retinal pigmentation typically observed significantly decreased RPE size and pigmentation (Baumann et al., 2016; Le et al., 2012). The RPE monolayer, embedding the outer segments of the photoreceptors has several important functions for the visual system such as nutrient and ion supply and retinol metabolism (Strauss, 2005). A stronger pigmentation might be related to a higher melanin content, which *per se* should have no negative effect on the larvae, since it is essential for an efficient light absorption. However, an exclusively beneficial role of melanin in RPE cells is still controversially discussed (Seagle et al., 2005) as evidence for a phototoxic effect of melanin in ROS photoproduction is also described (Boulton et al., 2001). Interestingly, alterations of the RPE were only observed in the LEWAF treatment group, although CEWAF is typically reported to induce overall stronger effects due to increased bioavailability of toxic oil components (e.g. Ramachandran et al., 2004). The reason for this cannot be identified within the current study since it might be related to the impact of the dispersant itself or an artefact due to histological section preparations. Hence, the role of the RPE should be addressed in more detail in future research.

#### Swimming behavior

Exposure to both, LEWAF and CEWAF induced a significant decrease in swimming activity of zebrafish embryos in response to the dark periods during the light/dark transition test (Figure 4, SI 1 Figure SI 2). Effects on swimming activity of fish at different life stages from early embryo to juveniles and adults induced by different types of crude oil or single and mixture PAHs have been described in several studies for different fish species. In line with the present findings, the main adverse effect observed in these studies was reduced swimming activity (Arukwe et al., 2008; Mager et al., 2014; Perrichon et al., 2014). However, increased swimming activity, greater mobility as well as anxiety-related alterations in shelter-seeking behavior has also been reported (Le Bihanic et al., 2014; Philibert et al., 2016; Vignet et al., 2014).

**Figure 4.**
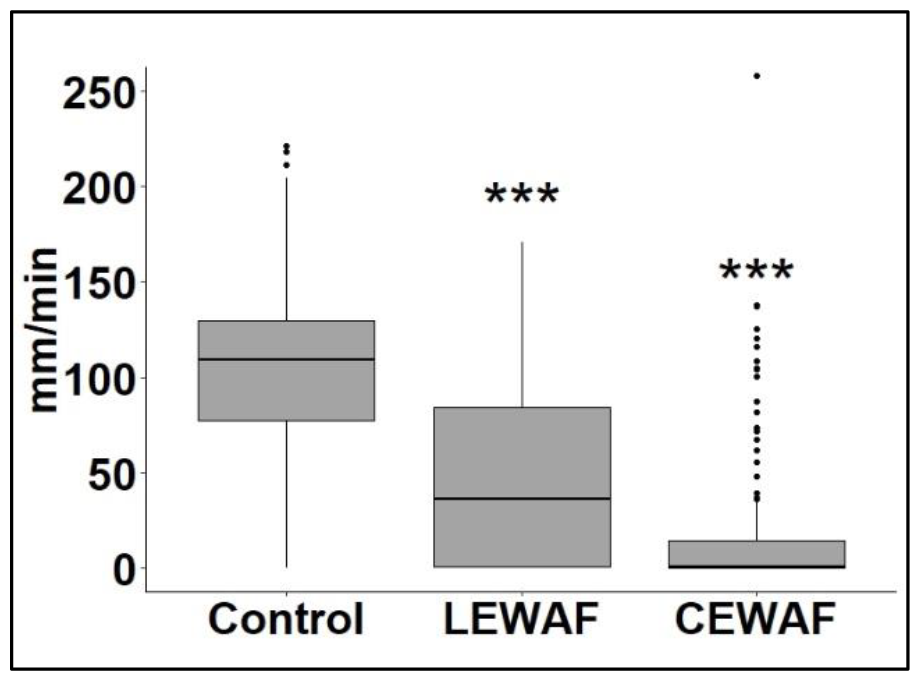
Swimming activity of 119 hpf zebrafish embryos exposed to WAF dilutions of a naphthenic North Sea crude oil and the dispersant Finasol OSR 51. Total distance moved by the embryos was pooled for min 26 for each treatment group. Results for the overall swimming distance were tested for normal distribution of the data using the Shapiro-Wilk test. Since the data was not normally distributed, Kruskall Wallis test with subsequent pairwise Wilcoxon rank sum test was used to test for significant differences between control and treatment groups. n = 4; **\*** = p value ≤ 0.05; **\*\*** = p value ≤ 0.01; **\*\*\*** = p value ≤ 0.001.

Effects on the swimming behavior of fish embryos is often used as a measure for developmental neurotoxicity (de Esch et al., 2012; Legradi et al., 2015, 2018; Perrichon et al., 2016; Selderslaghs et al., 2010). However, reduced swimming activity might not be exclusively related to neurotoxic effects but could rather be a secondary effect due to several other physiological issues. Several molecular and physiological effects have been proposed to be responsible for the reduced behavioral responsiveness in crude oil-exposed fish embryos. Exposure to AhR-agonists is, for example, known to result in the failure to inflate the swim bladder, which is proposed to affect behavior in fish embryos (Chatain, 1994; King Heiden et al., 2009; Marty et al., 1995). Other studies explained altered swimming patterns of fish due to reduced aerobic capacity resulting from cardiotoxicity (C. E. Hicken et al., 2011). Also, various sublethal effects of AhR-agonists and the resulting metabolic and energetic strain might lead to impaired swimming behavior (Correia et al., 2007). Importantly, some studies have related disturbances of normal functioning of the eyes to behavioral abnormalities (Kawaguchi et al., 2012; Magnuson et al., 2020; Philibert et al., 2021). A line of evidence exists that links eye development and the ability to display normal behavior in fish embryos which were exposed to crude oil or single AhR-agonists. Magnuson et al. (2020) showed that exposure to a weathered crude oil led to a decrease of the optomotor response of zebrafish embryos, demonstrating that the embryos’ ability to recognize moving objects was negatively affected. As we assessed the embryos’ swimming behavior in the light/dark transition test, we could evaluate their ability to differentiate between and react to different light intensities. The following section provides a detailed discussion about potential underlying molecular mechanisms linking eye function, morphological effects of the eye structure and behavior.

### 3.3. Linking results of functional annotation analysis to morphological and behavioral levels

#### General evaluation of transcriptional response

The statistical analysis of mRNA levels of control and LEWAF replicates revealed 101 genes that were dysregulated more than 1.5-fold compared to the control expression level (SI 2). Out of these, 17 were altered by more than 2-fold. Comparison of the control and CEWAF groups found a significantly different expression of 552 genes dysregulated more than 1.5-fold compared to control group expression levels (SI 3). For CEWAF, 106 genes were dysregulated more than 2-fold.

The comparison of the results obtained by qPCR vs RNAseq showed that fold-changes were in the same range of magnitude for all tested genes in both treatment groups (SI 1 Figure SI 3). Hence, it can be assumed that the detected fold-changes in the RNA sequencing analysis for significantly regulated genes were quantitatively in the same range of accuracy of PCR analysis techniques.

The functional annotation analysis identified terms belonging to various biological processes to be significantly (p-value < 0.05) dysregulated in crude oil WAF treated embryos. The analysis found 27 significantly enriched terms for LEWAF-treated embryos and 62 significantly enriched terms for CEWAF-treated embryos (SI 4 and 5). The most affected biological processes were related to xenobiotic biotransformation and visual stimuli. Dysregulation of genes involved in detoxification processes were among the most obviously affected, including genes regulated through the aryl hydrocarbon receptor (AhR) pathway (fold-change compared to control CEWAF: *cyp1a* (43.5), *ahrrb* (4.5); LEWAF: *cyp1a* (6.0), *ahrrb* (1.7)).

#### Eye development and function

Among the 20 most dysregulated genes in the LEWAF and CEWAF-treated groups were 3 genes known to be involved in the regulation of the development of the eyes (*rho, opn1sw1, rom1a*, all down-regulated) (SI 1 Table SI 4). The functional annotation analysis revealed that several pathways related to the development and function of specific compartments of the eyes were dysregulated (Figure 5, complete list of annotated genes SI 1 Table SI 5).

**Figure 5.**
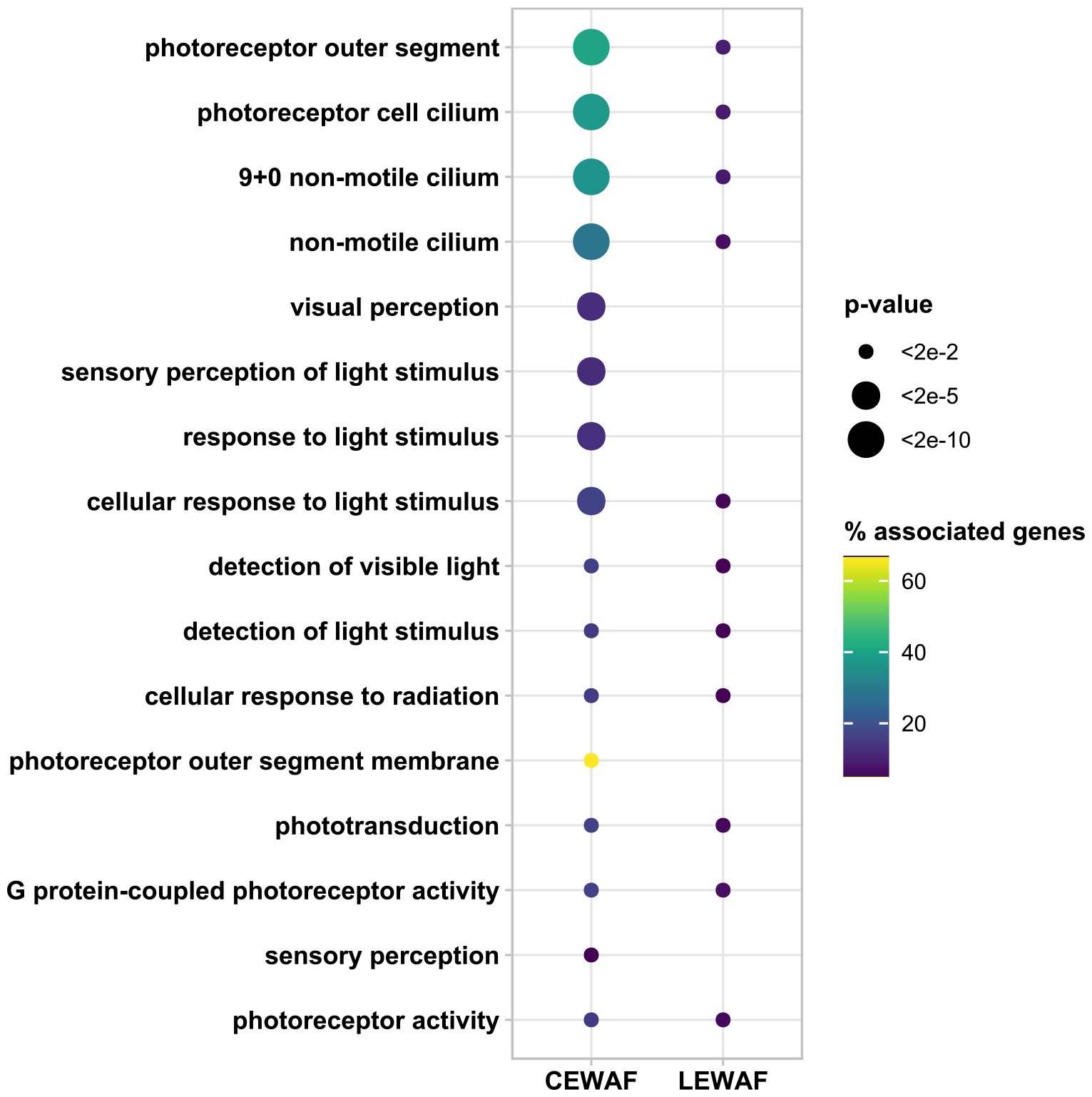
Pathways related to eye development and function significantly regulated in 119 hpf zebrafish embryos exposed to WAF dilutions of a naphthenic North Sea crude oil and the dispersant Finasol OSR 51. Zebrafish embryos were chronically exposed to WAF dilutions of crude oil only (LEWAF) and dispersed crude oil (CEWAF) at sublethal effect concentrations (EC10) until 119 hpf. The pathway enrichment analysis was performed in Cytoscape using the plug-in ClueGO. All dysregulated genes (fold-change > 1.5 compared to control) were included in the analysis. Significantly enriched (p-value< 0.05) pathways are displayed as scatterplots with node sizes indicating significance levels of the pathways (term p-values corrected for multiple testing with Bonferroni step down test) and node color indicating the percentage of associated genes that were matched to the displayed pathways.

While assigned pathways and terms regarding eye development and structure included particularly the photoreceptor layer (*photoreceptor outer segment, photoreceptor cell cilium* and *non-motile cilium*) the impaired function of the eye was highlighted by terms such as *phototransduction, detection of light stimulus, photoreceptor activity*. The evidence of impaired photoreceptors on transcriptional level directly links to the thinner photoreceptor layer from histological analyses.

The functional annotation analysis did not reveal dysregulation of specific terms that could explain the significantly reduced lens diameters in oil exposed embryos. However, the RNA-sequencing revealed that 30 genes encoding proteins of the crystallin family were downregulated 1.13-to 1.32-fold. Those proteins are the major structural and protective protein components of the vertebrate eye lens (Posner et al., 2008). A dysregulation of crystallins might indicate changes in the composition of the lens proteins, which has been shown to lead to the loss of lens transparency and partial or complete loss of vision (Bloemendal et al., 2004; Rao et al., 2011). It has to be considered that a conservative cutoff criterium (e.g. 1.5-fold change) has been discussed to not account for response variability (McCarthy and Smyth, 2009), probably leading to an underestimation of adverse effects and prohibiting the identification of affected pathways. Hence, a contribution of the lens in postulated impaired visual pathways needs to be addressed in more detail in future research.

The annotation terms *photoreceptor outer segment* and *photoreceptor cilium* (Go terms *cell cilium, (9+0) non-motile cilium*), significantly regulated in both treatment groups, directly link impaired structure with impaired function. The outer segment contains the overall phototransduction machinery, mainly driven by the visual pigments (Morris and Fadool, 2005). Opsins, the light-sensitive visual pigments of vertebrate photoreceptor cells, were among the most dysregulated genes in oil exposed fish. Different short-, medium- and long-wavelength opsins of cone photoreceptors were downregulated in LEWAF and CEWAF treatment groups, with the exception of *opn1mw2*, which was upregulated in both treatment groups (SI 2 and 3). In addition, the gene for the rod photoreceptor specific protein rhodopsin was also downregulated. Both cone and rod photoreceptors mediate zebrafish vision (Morris and Fadool, 2005), with rod photoreceptors particularly mediating the vision in dim light (Nathans, 1992) and cone photoreceptor cells taking over day-time vision (Raymond et al., 1995). Opsins are expressed from 50 hpf and the entire retina is laminated with rods and cones (Raymond et al., 1995) at 3 dpf. The signal transmission from photoreceptors to second order neurons becomes functional at the same time. Therefore, it can be assumed that the entire visual system is fully functional in 5 dpf zebrafish embryos used in the present study (Avanesov and Malicki, 2010). Importantly, the embryonal vision around this developmental stage is largely dominated by cone photoreceptors. The rod photoreceptor function is active at later developmental stages (around 15 dpf) (Bilotta et al., 2001; Clark, 1981; Morris and Fadool, 2005). Hence, the dysregulation of the opsins suggests that cone photoreceptor cell function in the zebrafish eyes was impaired by exposure to LEWAF and CEWAF. Additionally, many other genes encoding for photoreceptor disc membrane-associated proteins such as guanylyl cyclase, rhodopsin kinase or plasma membrane-associated ion exchanger proteins were significantly downregulated in oil exposed fish embryos (SI 2 and 3). These proteins are crucial for the functioning of photoreceptor cells to translate the light stimulus via hyperpolarization into an electric signal and also to recover from a light stimulus (Larhammar et al., 2009).

The downregulation of genes associated with the *photoreceptor cilium* also indicates an interrupted transport of proteins from the outer photoreceptor segment, containing the opsins, to the inner photoreceptor segment, since the cilium functions as the morphological bridge for the phototransduction cell signaling cascade (Goldberg et al., 2016). As reviewed by Mitchison and Valente (2017) the impairment of the protein traffic across the 9+0 cilium can have severe consequences for eye development and function.

In consequence of impaired phototransduction the visual perception might have been disturbed and could present an explanation for the lack of a normal response in the behavioral assay. Potentially, the lower expression of cone-specific genes important for proper photoreceptor function prevented the embryos in detecting the differences in light intensities during the light/dark transition test and hence embryos were not challenged to display increased swimming activities during the dark phases.

At this point it cannot be determined with certainty what the underlying molecular initiating events for the observed effects on the visual system were, and it is rather likely that these were based on a combination of multiple factors. However, some line of evidence indicate an AhR-related pathway, which is well known for cardiotoxic mode of action in oil exposed fish (Incardona, 2017). The AhR pathway was activated as expression levels of *cyp1a, ahrrb* and *ahr2* were significantly higher in comparison to the control group for both treatments (SI 2 and SI 3).

The activation of the AhR has been reported to affect retinol metabolism. Lie et al. (2019) showed the disruption of the retinoid acid signaling pathway by crude oil exposure in fish larvae, which can lead to abnormal eye development. However, retinol metabolism was not highlighted by functional annotation analysis even though genes associated with the retinol metabolism were impaired in CEWAF exposed fish embryos in the present study (*bco1l* 2.28-fold, *cyp1a* 44.51-fold, *cyp3a6* 51.82-fold regulation compared to control).

A direct connection between AhR responsive genes and normal eye development in fish has also been suggested by Aluru et al. (2014). In their study they found that a knockdown of the AhR repressor alpha (AHRRa) and beta (AHRRb) in zebrafish embryos affected the expression of genes involved in cone photoreceptor signaling, different opsins and crystallins. The authors suggested that distinct roles of AHRRa and AHRRb during zebrafish embryogenesis include eye development and function. However, since the knockdown of *ahrra* and *ahrrb* might have caused the failure of the AhR/AHRR regulatory loop it could not definitely be concluded which part of the gene cascade affected eye development. The activation of the AhR as a possible cause for the impairment of eye regulation was proposed by Huang et al. (2013). The authors found that exposure of zebrafish embryos to phenanthrene caused multiple morphological effects in the eyes, including reduction of thickness and loss of retina layers. The authors suggested that the repression of different transcription factors, such as *pax6*, caused by AhR up-regulation, lead to the observed effects on eye morphology. It has been shown that *pax6* is directly involved in the development of the vertebrate eye (Halder et al., 1995) and mutations of this gene is associated with malformations and diseases related to the eye (Hanson and Van Heyningen, 1995; Hill et al., 1991; Ton et al., 1991). In the present study both *pax6a* and *pax6b* were slightly but not significant downregulated in CEWAF treatment groups (*pax6a* -1.17; *pax6b* -1.17 (SI 3)) and LEWAF treatment groups (*pax6b* -1.11; *pax6a* not significantly regulated (SI 2)). Since this is only a slight alteration not even mirrored in a significant functional annotation, future studies will need to investigate what degree of dysregulation in *pax6* genes is sufficient to cause morphological effects in the eyes of exposed fish embryos. Additionally, AhR-independent pathways have to be investigated in future.

#### Circadian clock

Closely interlinked with pathways indicative for phototransduction is the circadian clock and circadian regulation. Genes associated with the circadian clock and annotated within phototransduction pathways were significantly regulated > 1.5-fold compared to the control group in CEWAF exposure (fold change in CEWAF exposure: *cry5*: 2.11, *nfiI3-6*: 2.22, *npas2*: 2.0, *per2*: 1.74, *cry1bb*: 1.55), while in LEWAF exposed embryos effects were less obvious (only *cry1bb* fold change >1.5). The mechanism driving the circadian rhythm in vertebrates can be divided into a positive and a negative feedback loop. During the positive feedback loop the CLOCK and ARNTL proteins drive the regulation of *cry* and *per* transcription. The CRY and PER proteins dimerize and inhibit the transcription of *clock* and *arntl* and thereby their own expression (Lowrey and Takahashi, 2004; McIntosh et al., 2010). The expression levels of the circadian clock regulator *per2* was significantly dysregulated, however, *clock* and *arntl* were not differentially expressed in CEWAF and LEWAF treatment groups (SI 2 and SI 3). The dysregulations of the clock regulator could suggest that the circadian regulation of the core clock or circadian gene regulation was affected at least in the CEWAF treatment.

The circadian rhythm of the behavior and physiology in mammals and teleost fish occurs in 24 h intervals and allows the organisms to adapt to their external environment and optimize metabolic processes in response to the day:night cycle (Ramasamy et al., 2019). The core clock can be entrained to external conditions and in mammals this mechanism is mediated mainly via the retina to the suprachiasmatic nucleus (SCN) (Ralph et al., 1990; Sack et al., 1992). In zebrafish, as in other organisms, the retina plays a distinctive role in the entrainment of the circadian rhythm as it expresses the majority of circadian clock genes (Ramasamy et al., 2019). The changes in transcription levels of circadian clock genes in zebrafish embryos exposed to LEWAF and CEWAF could be explained by the dysregulation of the retina-specific light sensitive opsins, since photoreceptors comminate the signals to the bipolar cells of the inner plexiform layer, which has recently been described to hold the majority of circadian clock genes (Ramasamy et al., 2019). If fewer functional opsins were present in the eyes of the embryos in the treatment group, this could have downstream effects on the entrainment and normal function of the circadian clock. In addition to the dysregulation of circadian genes through the function of the retina, the effects could have been mediated via the initial upregulation of transcription of *ahr2* that was observed in the present study. It has been suggested that AhR and ARNT can interact with circadian clock elements through promiscuous heterodimer formation, which could lead to the impairment of the circadian rhythm (Tischkau et al., 2011). This interaction of AhR signaling after exposure to AhR-ligands with core genes of the circadian clock has been demonstrated in mice (Tischkau et al., 2011), potentially directly through promiscuous interaction of the AhR and BMAL1 (brain and muscle ARNT-like) (Jaeger and Tischkau, 2016). However, effects of AhR-ligands on the circadian clock system of fish embryos have not been described so far. To obtain a better understanding of the circadian clock as potential target of AhR ligands in fish embryos and the potential consequences, future experiments should include the evaluation of effects on the circadian clock.

Furthermore, it has been shown that the locomotor activity of zebrafish embryos is controlled by the circadian clock (Cahill et al., 1998), and exposure to certain toxins can alter the circadian rhythm in fish, which can lead to abnormal behavior (Baganz et al., 1998). Hence, the potential disruption of the circadian clock in crude oil WAF exposed embryos could be a contributor to the detected reduced locomotor activity.

### 3.4. Effects of the chemical dispersant

The effects of the CEWAF on eye morphology, swimming behavior and gene expression in treated embryos were clearly stronger in comparison to LEWAF treated embryos. This discrepancy was not expected, since both LEWAF and CEWAF were tested at equipotent dilutions, which were based on lethal and sublethal effects in 5 dpf embryos determined in previous concentration-response experiments (Johann et al., 2020). Due to the limitations of the selected method for measurement of the soluble PAH fraction in CEWAF exposure medium, a comparison of the actual exposure concentrations of the target PAHs was not possible. However, it might be possible that concentrations of the compounds driving toxicity were elevated in CEWAF due to the presence of the dispersant. Additionally, the composition of the PAH fraction might have been different as has been suggested in one previous study using a different chemical analysis strategy in LEWAF and CEWAF (Esteban-Sánchez et al., 2021). Nonetheless, the elevated toxicity in CEWAFs highlights the controversial nature of the application of dispersants as an oil spill response measure in the environment. Dispersants are used to help protecting sensitive shoreline habitats (Lessard and DeMarco, 2000) because they facilitate the introduction of toxic compounds into the water column where they can be degraded by microbial processes. However, this process also makes these compounds more bioavailable for pelagic organisms, which are exposed to higher concentrations of the toxic fraction of crude oil until those substances are degraded, which is especially problematic for developing fish since they are very sensitive to the toxicity of PAHs. Overall, the present study did not investigate the dispersant itself since only a marginal contribution to toxicity in chemically dispersed crude oils in the tested concentration range was assumed based on previous studies (Johann et al., 2020) and the focus of the present study was on the mechanistic understanding of underlying effects.

## 4. Conclusion

A normally functioning visual system is crucial for the behavioral spectrum of fish. Failure of normal interactions between organisms and behavior like the search for food or predator avoidance potentially threatens the individuals’ life span and success in producing offspring. The present study showed that exposure to LEWAF and CEWAF caused an impairment of the visual system of developing zebrafish larvae across different levels of biological organization. This study contributes to the increasing body of literature that identifies oculotoxicity as an important adverse outcome in petroleum exposed fish embryos in addition to cardiotoxicity that was traditionally considered as the predominant adverse outcome. It can be highlighted that transcriptomic responses enabled the identification of potential novel pathways for crude oil mediated toxicity across levels of biological organisation. The present findings suggest that underlying mechanisms might be associated with the initial induction of the AhR-mediated pathways since there exist multiple studies suggesting a regulatory function of this receptor in development and function of the eye. In addition, the present study discussed the role of the circadian clock in oil toxicity as these types of effects have, to our knowledge, not been previously described in crude oil WAF-exposed fish embryos. Though first indications of dysregulated genes associated to circadian clock were observed, it could be concluded that more research is needed to fully understand the role in fish behavior and morphology. The present study also showed that adverse effects on sensitive life stages can occur in initially morphologically inconspicuous embryos. Comparable experimental setups with peak exposure and recovery phases should be repeated to fully understand the role of an impaired visual system as it is known that the teleost retina grows throughout its life and can regenerate from lesions. Hence, any adverse effects on the visual abilities of the fish embryos might be recovered later in life. The zebrafish model was used as a screening tool for specific underlying molecular effects that in future studies should be validated for environmentally relevant species.

Furthermore, future studies should investigate the impact of oil exposure on other sensory organs such as the lateral line and the olfactory system to fully understand potential mode of actions. The abundance of different pathways with potential severe outcomes that have been demonstrated in zebrafish embryos indicate that there exist different endpoints with great significance that are present after exposure to low concentrations of the bioavailable fraction of crude oil in the water column.

## Acknowledgements

This study has been carried out as part of the project ‘‘Integrated oil spill response actions and environmental effects — GRACE”, which is funded by the European Union’s Horizon 2020 research and innovation programme under grant agreement No 679266. This work was supported by the “Immunohistochemistry facility”, a core facility of the Interdisciplinary Center for Clinical Research (IZKF) Aachen within the Faculty of Medicine at RWTH Aachen University. The authors thank Dr. Lisa Baumann for her valuable insights on the histological analysis of zebrafish embryos. The authors would also like to kindly thank Noldus (Noldus Information Technology) and Nikon (Nikon GmbH) for their contribution to this study as partner of the Students Lab “Fascinating Environment” at Aachen Biology and Biotechnology (ABBT).

## Notes

### Competing Interest Statement

The authors have declared no competing interest.

## References

Aluru, N., Jenny, M.J., Hahn, M.E., 2014. Knockdown of a Zebrafish Aryl Hydrocarbon Receptor Repressor (AHRRa) Affects Expression of Genes Related to Photoreceptor Development and Hematopoiesis. Toxicol. Sci. 139, 381–395. 10.1093/toxsci/kfu052

Arukwe, A., Nordtug, T., Kortner, T.M., Mortensen, A.S., Brakstad, O.G., 2008. Modulation of steroidogenesis and xenobiotic biotransformation responses in zebrafish (Danio rerio) exposed to water-soluble fraction of crude oil. Environ. Res. 107, 362–370. 10.1016/j.envres.2008.02.009

Avanesov, A., Malicki, J., 2010. Chapter 6 - Analysis of the Retina in the Zebrafish Model, in: Detrich, H.W., Westerfield, M., Zon, L.I. (Eds.), Methods in Cell Biology, The Zebrafish: Cellular and Developmental Biology, Part A. Academic Press, pp. 153–204. 10.1016/B978-0-12-384892-5.00006-2

Baganz, D., Staaks, G., Steinberg, C., 1998. Impact of the cyanobacteria toxin, microcystin-lr on behaviour of zebrafish, danio rerio. Water Res. 32, 948–952. 10.1016/S0043-1354(97)00207-8

Baumann, L., Ros, A., Rehberger, K., Neuhauss, S.C.F., Segner, H., 2016. Thyroid disruption in zebrafish (Danio rerio) larvae: Different molecular response patterns lead to impaired eye development and visual functions. Aquat. Toxicol. 172, 44–55. 10.1016/j.aquatox.2015.12.015

Bilotta, J., Saszik, S., Sutherland, S.E., 2001. Rod contributions to the electroretinogram of the dark-adapted developing zebrafish. Dev. Dyn. 222, 564–570. 10.1002/dvdy.1188

Bindea, G., Mlecnik, B., Hackl, H., Charoentong, P., Tosolini, M., Kirilovsky, A., Fridman, W.-H., Pagès, F., Trajanoski, Z., Galon, J., 2009. ClueGO: a Cytoscape plug-in to decipher functionally grouped gene ontology and pathway annotation networks. Bioinformatics 25, 1091–1093. 10.1093/bioinformatics/btp101

Bloemendal, H., de Jong, W., Jaenicke, R., Lubsen, N.H., Slingsby, C., Tardieu, A., 2004. Ageing and vision: structure, stability and function of lens crystallins. Prog. Biophys. Mol. Biol. 86, 407–485. 10.1016/j.pbiomolbio.2003.11.012

Bornstein, J.M., Adams, J., Hollebone, B., King, T., Hodson, P.V., Brown, R.S., 2014. Effects-driven chemical fractionation of heavy fuel oil to isolate compounds toxic to trout embryos. Environ.Toxicol. Chem. 33, 814–824. 10.1002/etc.2492

Boulton, M., Różanowska, M., Różanowski, B., 2001. Retinal photodamage. J. Photochem. Photobiol. B, ESP Conference on Photoprotection 64, 144–161. 10.1016/S1011-1344(01)00227-5

Cahill, G.M., Hurd, M.W., Batchelor, M.M., 1998. Circadian rhythmicity in the locomotor activity of larval zebrafish. Neuroreport 9, 3445–3449. 10.1097/00001756-199810260-00020

Chatain, B., 1994. Abnormal swimbladder development and lordosis in sea bass (Dicentrarchus labrax) and sea bream (Sparus auratus). Aquaculture 119, 371–379. 10.1016/0044-8486(94)90301-8

Chen, J., Zhang, W., Wan, Z., Li, S., Huang, T., Fei, Y., 2019. Oil spills from global tankers: Status review and future governance. J. Clean. Prod. 227, 20–32. 10.1016/j.jclepro.2019.04.020

Clark, D.T., 1981. Visual responses in developing zebrafish (Ph. D. thesis). University of Oregon.

Correia, A.D., Gonçalves, R., Scholze, M., Ferreira, M., Henriques, M.A.-R., 2007. Biochemical and behavioral responses in gilthead seabream (Sparus aurata) to phenanthrene. J. Exp. Mar. Biol. Ecol. 347, 109–122. 10.1016/j.jembe.2007.03.015

Crowe, K.M., Newton, J.C., Kaltenboeck, B., Johnson, C., 2014. Oxidative stress responses of gulf killifish exposed to hydrocarbons from the Deepwater Horizon oil spill: Potential implications for aquatic food resources. Environ. Toxicol. Chem. 33, 370–374. 10.1002/etc.2427

de Esch, C., Slieker, R., Wolterbeek, A., Woutersen, R., de Groot, D., 2012. Zebrafish as potential model for developmental neurotoxicity testing: A mini review. Neurotoxicol. Teratol. 34, 545–553. 10.1016/j.ntt.2012.08.006

de Soysa, T.Y., Ulrich, A., Friedrich, T., Pite, D., Compton, S.L., Ok, D., Bernardos, R.L., Downes, G.B., Hsieh, S., Stein, R., Lagdameo, M.C., Halvorsen, K., Kesich, L.-R., Barresi, M.J., 2012. Macondo crude oil from the Deepwater Horizon oil spill disrupts specific developmental processes during zebrafish embryogenesis. BMC Biol. 10, 40. 10.1186/1741-7007-10-40

Dobin, A., Davis, C.A., Schlesinger, F., Drenkow, J., Zaleski, C., Jha, S., Batut, P., Chaisson, M., Gingeras, T.R., 2013. STAR: ultrafast universal RNA-seq aligner. Bioinformatics 29, 15–21. 10.1093/bioinformatics/bts635

Echols, B.S., Smith, A.J., Gardinali, P.R., Rand, G.M., 2015. Acute aquatic toxicity studies of Gulf of Mexico water samples collected following the Deepwater Horizon incident (May 12, 2010 to December 11, 2010). Chemosphere 120, 131–137. 10.1016/j.chemosphere.2014.06.048

Eckle, P., Burgherr, P., Michaux, E., 2012. Risk of Large Oil Spills: A Statistical Analysis in the Aftermath of Deepwater Horizon. Environ. Sci. Technol. 46, 13002–13008. 10.1021/es3029523

Esteban-Sánchez, A., Johann, S., Bilbao, D., Prieto, A., Hollert, H., Seiler, T.-B., Orbea, A., 2021. Multilevel responses of adult zebrafish to crude and chemically dispersed oil exposure. Environ. Sci. Eur. 33, 106. 10.1186/s12302-021-00545-4

EU, 2010. The European Parliament and the European Council Directive 2010/63/EU of the 22 September 2010 on the protection of animals used for scientific purposes. In: Off. J. Eur. Union (Hrsg.).

EU, 1986. Council Directive 86/609/EEC of 24 November 1986 on the approximation of laws, regulations and administrative provisions of the Member States regarding the protection of animals used for experimental and other scientific purposes. In: Off. J. Eur. Communities 29 L (Hrsg.).

Fadool, J.M., Dowling, J.E., 2008. Zebrafish: A model system for the study of eye genetics. Prog. Retin. Eye Res. 27, 89–110. 10.1016/j.preteyeres.2007.08.002

Fernald, R.D., 1985. Growth of the teleost eye: novel solutions to complex constraints. Environ. Biol. Fishes 13, 113–123. 10.1007/BF00002579

Fodrie, F.J., Jr., Heck, K.L., 2011. Response of Coastal Fishes to the Gulf of Mexico Oil Disaster. PLOS ONE 6, e21609. 10.1371/journal.pone.0021609

Gagnon, M.M., Holdway, D.A., 1999. Metabolic Enzyme Activities in Fish Gills as Biomarkers of Exposure to Petroleum Hydrocarbons. Ecotoxicol. Environ. Saf. 44, 92–99. 10.1006/eesa.1999.1804

Goldberg, A.F., Moritz, O.L., Williams, D.S., 2016. Molecular basis for photoreceptor outer segment architecture. Prog. Retin. Eye Res. 55, 52–81.

Gong, Y., Zhao, X., Cai, Z., O’Reilly, S.E., Hao, X., Zhao, D., 2014. A review of oil, dispersed oil and sediment interactions in the aquatic environment: Influence on the fate, transport and remediation of oil spills. Mar. Pollut. Bull. 79, 16–33. 10.1016/j.marpolbul.2013.12.024

Halder, G., Callaerts, P., Gehring, W.J., 1995. New perspectives on eye evolution. Curr. Opin. Genet. Dev. 5, 602–609. 10.1016/0959-437X(95)80029-8

Hanson, I., Van Heyningen, V.V., 1995. Pax6: more than meets the eye. Trends Genet. 11, 268–272. 10.1016/S0168-9525(00)89073-3

Hicken, Corinne E., Linbo, T.L., Baldwin, D.H., Willis, M.L., Myers, M.S., Holland, L., Larsen, M., Stekoll, M.S., Rice, S.D., Collier, T.K., Scholz, N.L., Incardona, J.P., 2011. Sublethal exposure to crude oil during embryonic development alters cardiac morphology and reduces aerobic capacity in adult fish. Proc. Natl. Acad. Sci. 108, 7086–7090. 10.1073/pnas.1019031108

Hicken, C. E., Linbo, T.L., Baldwin, D.H., Willis, M.L., Myers, M.S., Holland, L., Larsen, M., Stekoll, M.S., Rice, S.D., Collier, T.K., Scholz, N.L., Incardona, J.P., 2011. Sublethal exposure to crude oil during embryonic development alters cardiac morphology and reduces aerobic capacity in adult fish. Proc. Natl. Acad. Sci. 108, 7086–7090. 10.1073/pnas.1019031108

Hill, R.E., Favor, J., Hogan, B.L.M., Ton, C.C.T., Saunders, G.F., Hanson, I.M., Prosser, J., Jordan, T., Hastie, N.D., Heyningen, V. van, 1991. Mouse Small eye results from mutations in a paired-like homeobox-containing gene. Nature 354, 522–525. 10.1038/354522a0

Huang, L., Wang, C., Zhang, Y., Wu, M., Zuo, Z., 2013. Phenanthrene causes ocular developmental toxicity in zebrafish embryos and the possible mechanisms involved. J. Hazard. Mater. 261, 172–180. 10.1016/j.jhazmat.2013.07.030

Incardona, J.P., 2017. Molecular Mechanisms of Crude Oil Developmental Toxicity in Fish. Arch. Environ. Contam. Toxicol. 73, 19–32. 10.1007/s00244-017-0381-1

Incardona, J.P., Collier, T.K., Scholz, N.L., 2004. Defects in cardiac function precede morphological abnormalities in fish embryos exposed to polycyclic aromatic hydrocarbons. Toxicol. Appl. Pharmacol. 196, 191–205. 10.1016/j.taap.2003.11.026

ITOPF, 2022. Oil tanker spill statistics 2021. International Tanker Owners Pollution Federation Limited, London.

Jaeger, C., Tischkau, S.A., 2016. Role of Aryl Hydrocarbon Receptor in Circadian Clock Disruption and Metabolic Dysfunction. Environ. Health Insights 10, EHI.S38343. 10.4137/EHI.S38343

Johann, S., Nüßer, L., Goßen, M., Hollert, H., Seiler, T.B., 2020. Differences in biomarker and behavioral responses to native and chemically dispersed crude and refined fossil oils in zebrafish early life stages. Sci. Total Environ. 709, 136174. 10.1016/j.scitotenv.2019.136174

Johns, P.R., Easter, S.S., 1977. Growth of the adult goldfish eye. II. Increase in retinal cell number. J. Comp. Neurol. 176, 331–341. 10.1002/cne.901760303

Jørgensen, K.S., Kreutzer, A., Lehtonen, K.K., Kankaanpää, H., Rytkönen, J., Wegeberg, S., Gustavson, K., Fritt-Rasmussen, J., Truu, J., Kõuts, T., Lilover, M.-J., Seiler, T.-B., Hollert, H., Johann, S., Marigómez, I., Soto, M., Lekube, X., Jenssen, B.M., Ciesielski, T.M., Wilms, L.B., Högström, R., Pirneskoski, M., Virtanen, S., Forsman, B., Petrich, C., Phuong-Dang, N., Wang, F., 2019. The EU Horizon 2020 project GRACE: integrated oil spill response actions and environmental effects. Environ. Sci. Eur. 31, 44. 10.1186/s12302-019-0227-8

Khursigara, A.J., Perrichon, P., Martinez Bautista, N., Burggren, W.W., Esbaugh, A.J., 2017. Cardiac function and survival are affected by crude oil in larval red drum, Sciaenops ocellatus. Sci. Total Environ. 579, 797–804. 10.1016/j.scitotenv.2016.11.026

King Heiden, T.C., Spitsbergen, J., Heideman, W., Peterson, R.E., 2009. Persistent Adverse Effects on Health and Reproduction Caused by Exposure of Zebrafish to 2,3,7,8-Tetrachlorodibenzo-p-dioxin During Early Development and Gonad Differentiation. Toxicol. Sci. 109, 75–87. 10.1093/toxsci/kfp048

Larhammar, D., Nordström, K., Larsson, T.A., 2009. Evolution of vertebrate rod and cone phototransduction genes. Philos. Trans. R. Soc. B Biol. Sci. 364, 2867–2880. 10.1098/rstb.2009.0077

Le Bihanic, F., Clérandeau, C., Le Menach, K., Morin, B., Budzinski, H., Cousin, X., Cachot, J., 2014. Developmental toxicity of PAH mixtures in fish early life stages. Part II: adverse effects in Japanese medaka. Environ. Sci. Pollut. Res. 21, 13732–13743. 10.1007/s11356-014-2676-3

Le, H.-G.T., Dowling, J.E., Cameron, D.J., 2012. Early Retinoic acid deprivation in developing zebrafish results in microphthalmia. Vis. Neurosci. 29, 219–228. 10.1017/S0952523812000296

Legradi, J., el Abdellaoui, N., van Pomeren, M., Legler, J., 2015. Comparability of behavioural assays using zebrafish larvae to assess neurotoxicity. Environ. Sci. Pollut. Res. 22, 16277–16289. 10.1007/s11356-014-3805-8

Legradi, J.B., Di Paolo, C., Kraak, M.H.S., van der Geest, H.G., Schymanski, E.L., Williams, A.J., Dingemans, M.M.L., Massei, R., Brack, W., Cousin, X., Begout, M.-L., van der Oost, R., Carion, A., Suarez-Ulloa, V., Silvestre, F., Escher, B.I., Engwall, M., Nilén, G., Keiter, S.H., Pollet, D., Waldmann, P., Kienle, C., Werner, I., Haigis, A.-C., Knapen, D., Vergauwen, L., Spehr, M., Schulz, W., Busch, W., Leuthold, D., Scholz, S., vom Berg, C.M., Basu, N., Murphy, C.A., Lampert, A., Kuckelkorn, J., Grummt, T., Hollert, H., 2018. An ecotoxicological view on neurotoxicity assessment. Environ. Sci. Eur. 30, 46. 10.1186/s12302-018-0173-x

Lessard, R.R., DeMarco, G., 2000. The Significance of Oil Spill Dispersants. Spill Sci. Technol. Bull., 2000 Australia Oil Spill Response: 7th International Oil Spill 6, 59–68. 10.1016/S1353-2561(99)00061-4

Li, X., Xiong, D., Ju, Z., Xiong, Y., Ding, G., Liao, G., 2020. Phenotypic and transcriptomic consequences in zebrafish early-life stages following exposure to crude oil and chemical dispersant at sublethal concentrations. Sci. Total Environ. 143053. 10.1016/j.scitotenv.2020.143053

Liao, Y., Smyth, G.K., Shi, W., 2014. featureCounts: an efficient general purpose program for assigning sequence reads to genomic features. Bioinformatics 30, 923–930. 10.1093/bioinformatics/btt656

Lie, K.K., Meier, S., Sørhus, E., Edvardsen, R.B., Karlsen, Ø., Olsvik, P.A., 2019. Offshore Crude Oil Disrupts Retinoid Signaling and Eye Development in Larval Atlantic Haddock. Front. Mar. Sci. 6. 10.3389/fmars.2019.00368

Love, M.I., Huber, W., Anders, S., 2014. Moderated estimation of fold change and dispersion for RNA-seq data with DESeq2. Genome Biol. 15, 550. 10.1186/s13059-014-0550-8

Lowrey, P.L., Takahashi, J.S., 2004. Mammalian Circadian Biology: Elucidating Genome-Wide Levels of Temporal Organization. Annu. Rev. Genomics Hum. Genet. 5, 407–441. 10.1146/annurev.genom.5.061903.175925

Mager, E.M., Esbaugh, A.J., Stieglitz, J.D., Hoenig, R., Bodinier, C., Incardona, J.P., Scholz, N.L., Benetti, D.D., Grosell, M., 2014. Acute Embryonic or Juvenile Exposure to Deepwater Horizon Crude Oil Impairs the Swimming Performance of Mahi-Mahi (Coryphaena hippurus). Environ. Sci. Technol. 48, 7053–7061. 10.1021/es501628k

Magnuson, J.T., Bautista, N.M., Lucero, J., Lund, A.K., Xu, E.G., Schlenk, D., Burggren, W.W., Roberts, A.P., 2020. Exposure to Crude Oil Induces Retinal Apoptosis and Impairs Visual Function in Fish. Environ. Sci. Technol. 54, 2843–2850. 10.1021/acs.est.9b07658

Martin, J.D., Adams, J., Hollebone, B., King, T., Brown, R.S., Hodson, P.V., 2014. Chronic toxicity of heavy fuel oils to fish embryos using multiple exposure scenarios. Environ. Toxicol. Chem. 33, 677–687. 10.1002/etc.2486

Marty, G.D., Hinton, D.E., Jr, J.J.C., 1995. Notes: Oxygen Consumption by Larval Japanese Medaka with Inflated or Uninflated Swim Bladders. Trans. Am. Fish. Soc. 124, 623–627. 10.1577/1548-8659(1995)124<0623:NOCBLJ>2.3.CO;2

McCarthy, D.J., Smyth, G.K., 2009. Testing significance relative to a fold-change threshold is a TREAT. Bioinformatics 25, 765–771.

McIntosh, B.E., Hogenesch, J.B., Bradfield, C.A., 2010. Mammalian Per-Arnt-Sim Proteins in Environmental Adaptation. Annu. Rev. Physiol. 72, 625–645. 10.1146/annurev-physiol-021909-135922

Meador, J.P., Nahrgang, J., 2019. Characterizing Crude Oil Toxicity to Early-Life Stage Fish Based On a Complex Mixture: Are We Making Unsupported Assumptions? Environ. Sci. Technol. 53, 11080–11092. 10.1021/acs.est.9b02889

Mehinto, A.C., Martyniuk, C.J., Spade, D.J., Denslow, N.D., 2012. Applications for next-generation sequencing in fish ecotoxicogenomics. Front. Genet. 0. 10.3389/fgene.2012.00062

Mitchison, H.M., Valente, E.M., 2017. Motile and non‐motile cilia in human pathology: from functionto phenotypes. J. Pathol. 241, 294–309.

Morris, A.C., Fadool, J.M., 2005. Studying rod photoreceptor development in zebrafish. Physiol. Behav., Florida State University Special Issue 86, 306–313. 10.1016/j.physbeh.2005.08.020

Nahrgang, J., Camus, L., Gonzalez, P., Jönsson, M., Christiansen, J.S., Hop, H., 2010. Biomarker responses in polar cod (Boreogadus saida) exposed to dietary crude oil. Aquat. Toxicol. 96, 77–83. 10.1016/j.aquatox.2009.09.018

Nathans, J., 1992. Rhodopsin: structure, function, and genetics. Biochemistry 31, 4923–4931.

Pasparakis, C., Esbaugh, A.J., Burggren, W., Grosell, M., 2019. Physiological impacts of Deepwater Horizon oil on fish. Comp. Biochem. Physiol. Part C Toxicol. Pharmacol. 224, 108558. 10.1016/j.cbpc.2019.06.002

Perrichon, P., Le Bihanic, F., Bustamante, P., Le Menach, K., Budzinski, H., Cachot, J., Cousin, X., 2014. Influence of sediment composition on PAH toxicity using zebrafish (Danio rerio) and Japanese medaka (Oryzias latipes) embryo-larval assays. Environ. Sci. Pollut. Res. 21, 13703–13719. 10.1007/s11356-014-3502-7

Perrichon, P., Le Menach, K., Akcha, F., Cachot, J., Budzinski, H., Bustamante, P., 2016. Toxicity assessment of water-accommodated fractions from two different oils using a zebrafish (Danio rerio) embryo-larval bioassay with a multilevel approach. Sci. Total Environ. 568, 952–966. 10.1016/j.scitotenv.2016.04.186

Peterson, C.H., Rice, S.D., Short, J.W., Esler, D., Bodkin, J.L., Ballachey, B.E., Irons, D.B., 2003. Long-Term Ecosystem Response to the Exxon Valdez Oil Spill. Science 302, 2082–2086. 10.1126/science.1084282

Pfaffl, M.W., 2001. A new mathematical model for relative quantification in real-time RT–PCR. Nucleic Acids Res. 29, e45–e45. 10.1093/nar/29.9.e45

Philibert, D.A., Lyons, D.D., Tierney, K.B., 2021. Comparing the effects of unconventional and conventional crude oil exposures on zebrafish and their progeny using behavioral and genetic markers. Sci. Total Environ. 770, 144745. 10.1016/j.scitotenv.2020.144745

Philibert, D.A., Philibert, C.P., Lewis, C., Tierney, K.B., 2016. Comparison of Diluted Bitumen (Dilbit) and Conventional Crude Oil Toxicity to Developing Zebrafish. Environ. Sci. Technol. 50, 6091–6098. 10.1021/acs.est.6b00949

Posner, M., Hawke, M., LaCava, C., Prince, C.J., Bellanco, N.R., Corbin, R.W., 2008. A proteome map of the zebrafish (Danio rerio) lens reveals similarities between zebrafish and mammalian crystallin expression. Mol. Vis. 14, 806–814.

Potter, D.W., Pawliszyn, Janusz., 1994. Rapid determination of polyaromatic hydrocarbons and polychlorinated biphenyls in water using solid-phase microextraction and GC/MS. Environ. Sci. Technol. 28, 298–305. 10.1021/es00051a017

Prince, R.C., 2015. Oil Spill Dispersants: Boon or Bane? Environ. Sci. Technol. 49, 6376–6384. 10.1021/acs.est.5b00961

Ralph, M.R., Foster, R.G., Davis, F.C., Menaker, M., 1990. Transplanted suprachiasmatic nucleus determines circadian period. Science 247, 975–978. 10.1126/science.2305266

Ramachandran, S.D., Hodson, P.V., Khan, C.W., Lee, K., 2004. Oil dispersant increases PAH uptake by fish exposed to crude oil. Ecotoxicol. Environ. Saf. 59, 300–308. 10.1016/j.ecoenv.2003.08.018

Ramasamy, S., Sharma, S., Iyengar, B.R., Vellarikkal, S.K., Sivasubbu, S., Maiti, S., Pillai, B., 2019. Identification of novel circadian transcripts in the zebrafish retina. J. Exp. Biol. 222. 10.1242/jeb.192195

Rao, G.N., Khanna, R., Payal, A., 2011. The global burden of cataract. Curr. Opin. Ophthalmol. 22, 4–9. 10.1097/ICU.0b013e3283414fc8

Raymond, P.A., Barthel, L.K., Curran, G.A., 1995. Developmental patterning of rod and cone photoreceptors in embryonic zebrafish. J. Comp. Neurol. 359, 537–550. 10.1002/cne.903590403

Reddy, C.M., Eglinton, T.I., Hounshell, A., White, H.K., Xu, L., Gaines, R.B., Frysinger, G.S., 2002. The West Falmouth Oil Spill after Thirty Years: The Persistence of Petroleum Hydrocarbons in Marsh Sediments. Environ. Sci. Technol. 36, 4754–4760. 10.1021/es020656n

Sack, R.L., Lewy, A.J., Blood, M.L., Keith, L.D., Nakagawa, H., 1992. Circadian rhythm abnormalities in totally blind people: incidence and clinical significance. J. Clin. Endocrinol. Metab. 75, 127–134. 10.1210/jcem.75.1.1619000

Seagle, B.-L.L., Rezai, K.A., Kobori, Y., Gasyna, E.M., Rezaei, K.A., Norris, J.R., 2005. Melanin photoprotection in the human retinal pigment epithelium and its correlation with light-induced cell apoptosis. Proc. Natl. Acad. Sci. 102, 8978–8983. 10.1073/pnas.0501971102

Selderslaghs, I.W.T., Hooyberghs, J., De Coen, W., Witters, H.E., 2010. Locomotor activity in zebrafish embryos: A new method to assess developmental neurotoxicity. Neurotoxicol. Teratol. 32, 460–471. 10.1016/j.ntt.2010.03.002

Shannon, P., Markiel, A., Ozier, O., Baliga, N.S., Wang, J.T., Ramage, D., Amin, N., Schwikowski, B., Ideker, T., 2003. Cytoscape: a software environment for integrated models of biomolecular interaction networks. Genome Res. 13, 2498–2504. 10.1101/gr.1239303

Short, J.W., Lindeberg, M.R., Harris, P.M., Maselko, J.M., Pella, J.J., Rice, S.D., 2004. Estimate of Oil Persisting on the Beaches of Prince William Sound 12 Years after the Exxon Valdez Oil Spill. Environ. Sci. Technol. 38, 19–25. 10.1021/es0348694

Strähle, U., Scholz, S., Geisler, R., Greiner, P., Hollert, H., Rastegar, S., Schumacher, A., Selderslaghs, I., Weiss, C., Witters, H., Braunbeck, T., 2012. Zebrafish embryos as an alternative to animal experiments—A commentary on the definition of the onset of protected life stages in animal welfare regulations. Reprod. Toxicol., Zebrafish Teratogenesis 33, 128–132. 10.1016/j.reprotox.2011.06.121

Strauss, O., 2005. The Retinal Pigment Epithelium in Visual Function. Physiol. Rev. 85, 845–881. 10.1152/physrev.00021.2004

Tischkau, S.A., Jaeger, C.D., Krager, S.L., 2011. Circadian clock disruption in the mouse ovary in response to 2,3,7,8-tetrachlorodibenzo-p-dioxin. Toxicol. Lett. 201, 116–122. 10.1016/j.toxlet.2010.12.013

Ton, C.C.T., Hirvonen, H., Miwa, H., Weil, M.M., Monaghan, P., Jordan, T., van Heyningen, V., Hastie, N.D., Meijers-Heijboer, H., Drechsler, M., Royer-Pokora, B., Collins, F., Swaroop, A., Strong, L.C., Saunders, G.F., 1991. Positional cloning and characterization of a paired box- and homeobox-containing gene from the aniridia region. Cell 67, 1059–1074. 10.1016/0092-8674(91)90284-6

Tronczyński, J., Munschy, C., Héas-Moisan, K., Guiot, N., Truquet, I., Olivier, N., Men, S., Furaut, A., 2004. Contamination of the Bay of Biscay by polycyclic aromatic hydrocarbons (PAHs) following the T/V “Erika” oil spill. Aquat. Living Resour. 17, 243–259. 10.1051/alr:2004042

Vignet, C., Le Menach, K., Lyphout, L., Guionnet, T., Frère, L., Leguay, D., Budzinski, H., Cousin, X., Bégout, M.-L., 2014. Chronic dietary exposure to pyrolytic and petrogenic mixtures of PAHs causes physiological disruption in zebrafish—part II: behavior. Environ. Sci. Pollut. Res. 21, 13818–13832. 10.1007/s11356-014-2762-6

Vihtelic, T.S., Hyde, D.R., 2000. Light-induced rod and cone cell death and regeneration in the adult albino zebrafish (Danio rerio) retina. J. Neurobiol. 44, 289–307. 10.1002/1097-4695(20000905)44:3<289::AID-NEU1>3.0.CO;2-H

Wagle, P., Nikolić, M., Frommolt, P., 2015. QuickNGS elevates Next-Generation Sequencing data analysis to a new level of automation. BMC Genomics 16, 487. 10.1186/s12864-015-1695-x

Xu, E.G., Mager, E.M., Grosell, M., Pasparakis, C., Schlenker, L.S., Stieglitz, J.D., Benetti, D., Hazard, E.S., Courtney, S.M., Diamante, G., Freitas, J., Hardiman, G., Schlenk, D., 2016. Time- and Oil-Dependent Transcriptomic and Physiological Responses to Deepwater Horizon Oil in Mahi-Mahi (Coryphaena hippurus) Embryos and Larvae. Environ. Sci. Technol. 50, 7842–7851. 10.1021/acs.est.6b02205

