## Supplementary material for "Zebrafish eye development and function is affected by crude oil exposure – line of evidence from underlying molecular effects to behavioral consequences": File SI 1

***Table SI 1 Complete list of target compounds for chemical analysis of LEWAF exposure medium**. 18 PAHs were analyzed in the LEWAF exposure medium. Out of the 18 PAHs three were detected (D), the remaining 15 PAHs were not detected (N.D.) in the samples.*

| Target compound | D = detected; N.D. = not detected |
|---|---|
| Naphthalene | D |
| Fluorene | D |
| Phenathrene | D |
| Anthracene | N.D. |
| Fluoranthene | N.D. |
| Pyrene | N.D. |
| 11hbenzo[a]fluorene | N.D. |
| 11hbenzo[b]fluorene | N.D. |
| Benzo[a]anthracene | N.D. |
| Chrysene | N.D. |
| Benzo[b]fluoranthene | N.D. |
| Benzo[k]fluoranthene | N.D. |
| Benzo[a]pyrene | N.D. |
| Benzo[e]pyrene | N.D. |
| Indeno[1,2,3 cd]pyrene | N.D. |
| Dibenz[a,h]anthracen | N.D. |
| Benzo[ghi]perylene | N.D. |
| Dibenzo[a,e]pyren | N.D. |

***Table SI 2*** *Primer pair sequences, accession numbers and annealing temperatures used for Illumina sequencing validation via RT-qPCR. Specificity of all primers was confirmed by DNA oligo sequencing and by amplicon separation on 1% Agarose gels with DNA size standard (Quick load ladder N0469S, BioLabs).*

| gene | transcript ID | primer pair | (Tm) [°C] |
|---|---|---|---|
| *cyp1a* | NM_131879 | fw: TGCGAAGACCGAAAACTGGA | 60 |
| | | rev: TCGAAACCGGCTCCGAATAG | |
| *cyp1b1* | NM_001045256 | fw: TGGATCATCCTGCTACTTGTCA | 62 |
| | | rev: TCCACTACCCTGTCCACGTC | |
| *cyp1c1* | NM_001020610 | fw: GACTGAGTGCTGATGGACGA | 61 |
| | | rev: CACAGAGCGCAGATGACATT | |
| *ahrrb* | NM_001033920 | fw: ACCTGTGCAGAAACAGAAACC | 61 |
| | | rev: GCTCTGCATTCAGACGGTCA | |
| *sult6b1* | NM_214686 | fw: CCCTGACAACATGCCTGCTA | 60 |
| | | rev: GAGCCCCAACTAACATCACCA | |
| *nfe2l2a* | NM_182889 | fw: ACTCCCAGAGTTGCAGCAGT | 60 |
| | | rev: ACTTCTGTTTGAGCCGAGCC | |
| *rho* | NM_131084 | fw: ACTTCCGTTTCGGGGAGAAC | 60 |
| | | rev: GGAGTGCGGGTGTAGTAGTC | |
| *opn1sw1* | NM_131319 | fw: ATGGTCCTTGGCTGTTCTGG | 60 |
| | | rev: CCTCGGGAATGTATCTGCTCC | |
| opn1mw2 | NM_182891 | fw: TTGGCTGGTCCCGATACATCC | 60 |
| | | rev: CAGGAACGCAGAAATGACAGC | |
| eef1a1 | NM_131263 | fw: GTGGTAATGTGGCTGGAGAC | 60 |
| | | rev: TGTGAGCAGTGTGGCAATC | |

***Table SI 3 Chemical analysis of the LEWAF exposure medium.*** *PAHs were measured each day in old and fresh exposure medium (12.5 % of stock (LEWAF 1:400 dilution)). Out of 18 target PAHs, naphthalene, Fluorene and Phenanthrene were detected in the exposure medium. The concentrations of naphthalene and fluorene in the fresh medium on day 2 were outside the calibration range and could therefore not be quantified.*

| Day | Naphthalene (µg/L) | Fluorene (µg/L) | Phenanthrene (µg/L) |
|---|---|---|---|
| 1 (fresh medium) | 26.18 ± 8.43 | 0.41 ± 0.05 | 0.32 ± 0.03 |
| 2 (old medium) | 21.55 ± 5.56 | 0.28 ± 0.01 | 0.24 ± 0.02 |
| 2 (fresh medium) | no calibration | no calibration | 0.30 ± 0.02 |
| 3 (old medium) | 17.06 ± 6.50 | 0.33 ± 0.04 | 0.25 ± 0.02 |
| 3 (fresh medium) | 17.91 ± 4.16 | 0.41 ± 0.03 | 0.34 ±0.02 |
| 4 (old medium) | 19.02 ± 2.84 | 0.28 ± 0.02 | 0.25 ± 0.02 |
| 4 (fresh medium) | 33.16 ± 5.98 | 0.46 ± 0.07 | 0.40 ± 0.05 |
| 5 (old medium) | 24.97 ± 4.05 | 0.28 ± 0.04 | 0.19 ± 0.01 |
| 5 (fresh medium) | 29.35 ± 4.70 | 0.39 ± 0.04 | 0.31 ± 0.01 |
| 6 (old medium) | 23.14 ± 3.77 | 0.29 ± 0.05 | 0.16 ± 0.03 |

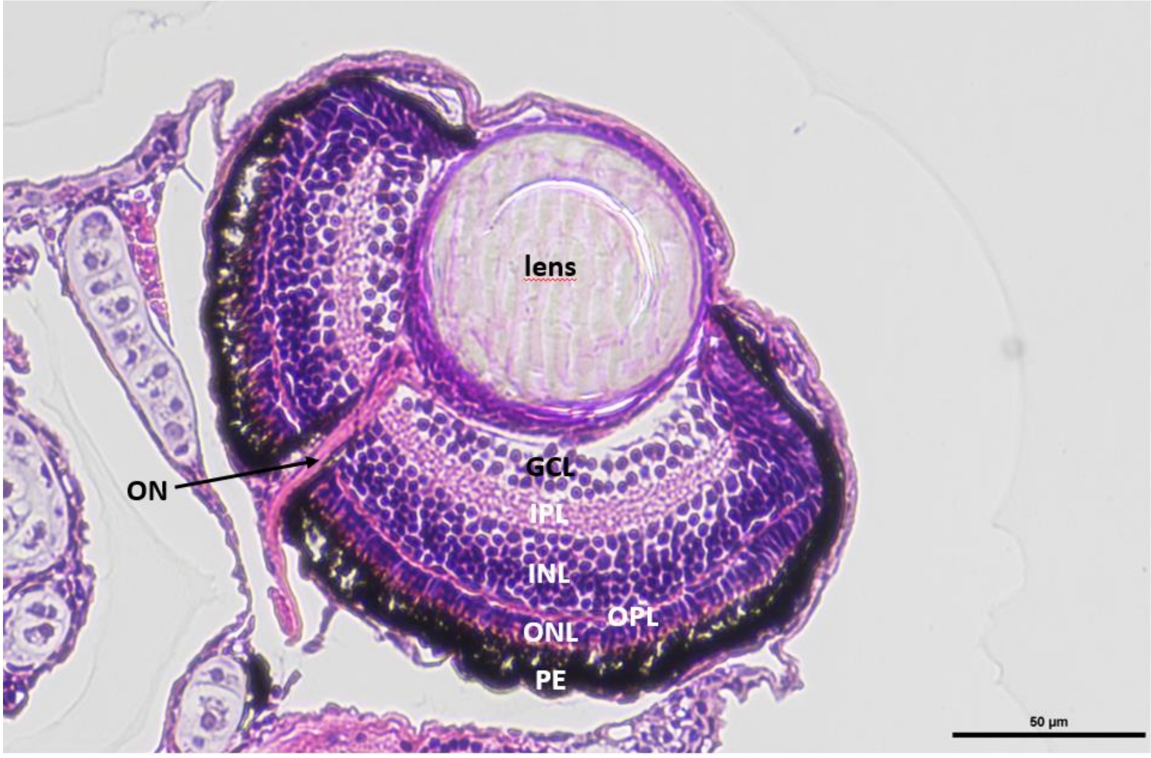


***Figure SI 1 Exemplary graph for the identity and location of the different layers of the zebrafish embryo retina****. Histological cut of the eye of a 119 hpf zebrafish embryo. ON = Optical nerve; GCL = Ganglion cell layer; IPL = Inner plexiform layer; INL = Inner nuclear layer; OPL = Outer plexiform layer; ONL = Outer nuclear layer; PE = Pigment epithelium.*

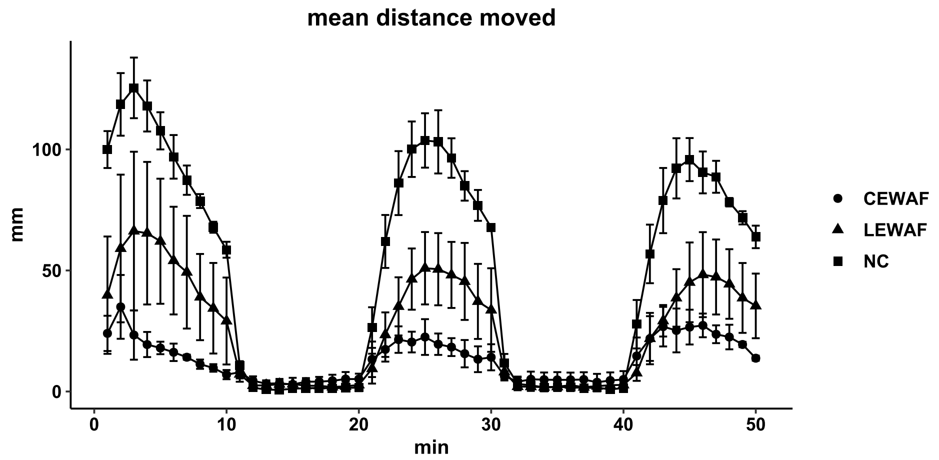


***Figure SI 2 Swimming activity during the light/dark transition test.*** Zebrafish embryos were chronically exposed to WAF dilutions of crude oil only (LEWAF) and dispersed crude oil (CEWAF) at sublethal effect concentrations (EC10) until 119 hpf. Swimming activity was pooled as mean distance moved (mm) in 1 min time bins for 4 replicates. n treatments was at least 28 individuals and maximal 30 individuals per replicate.

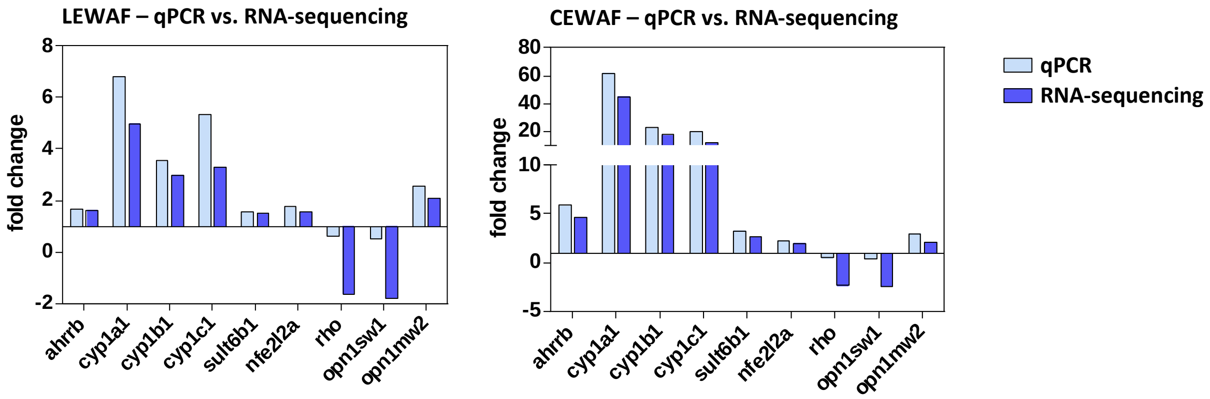


***Figure SI 3 Comparison of the transcript abundance of selected genes for LEWAF and CEWAF treatments between RNA-sequencing and qPCR results in 119 hpf zebrafish embryos.*** *Zebrafish embryos were chronically exposed to WAF dilutions of crude oil only (LEWAF) and dispersed crude oil (CEWAF) at sublethal effect concentrations (EC10) until 119 hpf. Effects levels of gene regulation were measured using whole transcriptome RNA-sequencing. The expression levels of selected genes were analyzed in subsamples using quantitative real-time PCR (qPCR) for validation of the RNA-sequencing data.*

***Table SI 4 List of 20 most dysregulated genes***. *Zebrafish embryos were chronically exposed to WAF dilutions of crude oil only (LEWAF) and dispersed crude oil (CEWAF) at sublethal effect concentrations (EC10) until 119 hpf. Effect levels of gene regulation were measured using whole transcriptome RNA-sequencing. In this list were only genes included that already have described functions and a gene symbol.*

| **LEWAF** | | **CEWAF** | |
|---|---|---|---|
| gene symbol | fold change relative to control | gene symbol | fold change relative to control |
| cyp1a | 6,0 | cyp1a | 43,5 |
| cyp1c1 | 4,0 | cyp1b1 | 18,1 |
| cyp1b1 | 3,1 | cyp1c1 | 14,0 |
| cyp2k20 | 2,3 | fgf7 | 6,1 |
| caspbl | 2,3 | ahrrb | 4,5 |
| abcb5 | 2,3 | prss60.3 | 3,8 |
| ugt5a1 | 2,1 | foxq1a | 3,4 |
| vwa10.1 | 2,1 | caspbl | 3,3 |
| abcc12 | 1,9 | gna15.2 | 3,2 |
| cyp2x10.2 | 1,9 | vwa10.1 | 3,2 |
| pdha1b | -1,5 | opn1sw1 | -2,2 |
| gngt1 | -1,5 | rom1a | -2,2 |
| rom1a | -1,5 | gngt1 | -2,3 |
| gnat1 | -1,5 | rho | -2,3 |
| rho | -1,5 | nr1d4b | -2,3 |
| opn1sw1 | -1,6 | gnat1 | -2,3 |
| nr2e3 | -1,6 | cyt1l | -2,4 |
| krt15 | -1,7 | nr2e3 | -2,4 |
| guca1a | -1,7 | guca1a | -2,5 |
| gck | -2,1 | gck | -2,7 |

***Table SI 5 Genes symbols and fold-changes for all pathways related to eye development and function.*** *Zebrafish embryos were chronically exposed to WAF dilutions of crude oil only (LEWAF) and dispersed crude oil (CEWAF) at sublethal effect concentrations (EC10) until 119 hpf. The pathway enrichment analysis was performed in Cytoscape using the plug-in ClueGO. All dysregulated genes (fold-change > 1.5 compared to control) were included in the analysis. Displayed is the term p-value corrected for multiple testing with Bonferroni step down test and all gene symbols with their measured expression value.*

| **Pathway** | **Corrected term p-value** | **Treatment** | **Data base** | **Associated genes with according fold change** |
|---|---|---|---|---|
| photoreceptor outer segment | 7,90E-16 | CEWAF | GO_CellularComponent-EBI-UniProt-GOA-ACAP-ARAP_16.06.2023_00h00 | gnat2, guca1b, nxnl1, opn1lw2, opn1mw1, opn1mw2, opn1sw1, opn1sw2, pdca, pde6a, pde6b, pde6ga, pde6gb, rho, rhol, saga |
| photoreceptor outer segment | 1,58E-04 | LEWAF | GO_CellularComponent-EBI-UniProt-GOA-ACAP-ARAP_16.06.2023_00h00 | opn1mw2, opn1sw1, pde6ga, rho |
| photoreceptor cell cilium | 3,30E-15 | CEWAF | GO_CellularComponent-EBI-UniProt-GOA-ACAP-ARAP_16.06.2023_00h00 | gnat2, guca1b, nxnl1, opn1lw2, opn1mw1, opn1mw2, opn1sw1, opn1sw2, pdca, pde6a, pde6b, pde6ga, pde6gb, rho, rhol, saga |
| photoreceptor cell cilium | 1,99E-04 | LEWAF | GO_CellularComponent-EBI-UniProt-GOA-ACAP-ARAP_16.06.2023_00h00 | opn1mw2, opn1sw1, pde6ga, rho |
| 9+0 non-motile cilium | 5,14E-15 | CEWAF | GO_CellularComponent-EBI-UniProt-GOA-ACAP-ARAP_16.06.2023_00h00 | gnat2, guca1b, nxnl1, opn1lw2, opn1mw1, opn1mw2, opn1sw1, opn1sw2, pdca, pde6a, pde6b, pde6ga, pde6gb, rho, rhol, saga |
| 9+0 non-motile cilium | 2,04E-04 | LEWAF | GO_CellularComponent-EBI-UniProt-GOA-ACAP-ARAP_16.06.2023_00h00 | opn1mw2, opn1sw1, pde6ga, rho |
| non-motile cilium | 6,42E-13 | CEWAF | GO_CellularComponent-EBI-UniProt-GOA-ACAP-ARAP_16.06.2023_00h00 | gnat2, guca1b, nxnl1, opn1lw2, opn1mw1, opn1mw2, opn1sw1, opn1sw2, pdca, pde6a, pde6b, pde6ga, pde6gb, rho, rhol, saga |
| non-motile cilium | 5,41E-04 | LEWAF | GO_CellularComponent-EBI-UniProt-GOA-ACAP-ARAP_16.06.2023_00h00 | opn1mw2, opn1sw1, pde6ga, rho |
| visual perception | 2,79E-09 | CEWAF | GO_BiologicalProcess-EBI-UniProt-GOA-ACAP-ARAP_16.06.2023_00h00 | gnat2, grk1a, guca1b, gucy2f, kera, opn1lw2, opn1mw1, opn1mw2, opn1sw1, opn1sw2, pdca, pde6ga, pde6gb, prph2a, prph2la, rgs9a, rho, rhol, rom1a, rom1b |
| sensory perception of light stimulus | 3,12E-09 | CEWAF | GO_BiologicalProcess-EBI-UniProt-GOA-ACAP-ARAP_16.06.2023_00h00 | gnat2, grk1a, guca1b, gucy2f, kera, opn1lw2, opn1mw1, opn1mw2, opn1sw1, opn1sw2, pdca, pde6ga, pde6gb, prph2a, prph2la, rgs9a, rho, rhol, rom1a, rom1b |

| | | | | |
|---|---|---|---|---|
| response to light stimulus | 1,19E-08 | CEWAF | GO_BiologicalProcess-EBI-UniProt-GOA-ACAP-ARAP_16.06.2023_00h00 | arl3l2, cry5, gnat1, gnat2, grk1a, grk1b, npas2, opn1lw2, opn1mw1, opn1mw2, opn1sw1, opn1sw2, per2, rcvrna, rho, rhol, si:ch1073-390k14.1, tefa |
| cellular response to light stimulus | 8,26E-05 | CEWAF | GO_BiologicalProcess-EBI-UniProt-GOA-ACAP-ARAP_16.06.2023_00h00 | grk1a, opn1lw2, opn1mw1, opn1mw2, opn1sw1, opn1sw2, rho, rhol, tefa |
| cellular response to light stimulus | 4,07E-03 | LEWAF | GO_BiologicalProcess-EBI-UniProt-GOA-ACAP-ARAP_16.06.2023_00h00 | opn1mw2, opn1sw1, rho |
| detection of visible light | 1,34E-04 | CEWAF | GO_BiologicalProcess-EBI-UniProt-GOA-ACAP-ARAP_16.06.2023_00h00 | gnat2, grk1a, opn1lw2, opn1mw1, opn1mw2, opn1sw1, opn1sw2, rho, rhol |
| detection of visible light | 3,84E-03 | LEWAF | GO_BiologicalProcess-EBI-UniProt-GOA-ACAP-ARAP_16.06.2023_00h00 | opn1mw2, opn1sw1, rho |
| detection of light stimulus | 2,11E-04 | CEWAF | GO_BiologicalProcess-EBI-UniProt-GOA-ACAP-ARAP_16.06.2023_00h00 | gnat2, grk1a, opn1lw2, opn1mw1, opn1mw2, opn1sw1, opn1sw2, rho, rhol |
| detection of light stimulus | 3,36E-03 | LEWAF | GO_BiologicalProcess-EBI-UniProt-GOA-ACAP-ARAP_16.06.2023_00h00 | opn1mw2, opn1sw1, rho |
| cellular response to radiation | 2,44E-04 | CEWAF | GO_BiologicalProcess-EBI-UniProt-GOA-ACAP-ARAP_16.06.2023_00h00 | grk1a, opn1lw2, opn1mw1, opn1mw2, opn1sw1, opn1sw2, rho, rhol, tefa |
| cellular response to radiation | 2,35E-03 | LEWAF | GO_BiologicalProcess-EBI-UniProt-GOA-ACAP-ARAP_16.06.2023_00h00 | opn1mw2, opn1sw1, rho |
| photoreceptor outer segment membrane | 3,11E-04 | CEWAF | GO_CellularComponent-EBI-UniProt-GOA-ACAP-ARAP_16.06.2023_00h00 | pde6a, pde6b, pde6ga, pde6gb |
| phototransduction | 4,68E-04 | CEWAF | GO_BiologicalProcess-EBI-UniProt-GOA-ACAP-ARAP_16.06.2023_00h00 | grk1a, opn1lw2, opn1mw1, opn1mw2, opn1sw1, opn1sw2, rho, rhol |
| phototransduction | 3,86E-03 | LEWAF | GO_BiologicalProcess-EBI-UniProt-GOA-ACAP-ARAP_16.06.2023_00h00 | opn1mw2, opn1sw1, rho |

| | | | | |
|---|---|---|---|---|
| G protein-couples photoreceptor activity | 1,94E-03 | CEWAF | GO_BiologicalProcess-EBI-UniProt-GOA-ACAP-ARAP_16.06.2023_00h00 | opn1lw2, opn1mw1, opn1mw2, opn1sw1, opn1sw2, rho, rhol |
| G protein-couples photoreceptor activity | 3,47E-03 | LEWAF | GO_BiologicalProcess-EBI-UniProt-GOA-ACAP-ARAP_16.06.2023_00h00 | opn1mw2, opn1sw1, rho |
| Sensory perception | 2,64E-03 | CEWAF | GO_BiologicalProcess-EBI-UniProt-GOA-ACAP-ARAP_16.06.2023_00h00 | adcy1b, gnat2, grk1a, guca1b, gucy2f, kera, opn1lw2, opn1mw1, opn1mw2, opn1sw1, opn1sw2, pdca, pde6ga, pde6gb, prph2a, prph2la, rgs9a, rho, rhol, rom1a, rom1b |
| photoreceptor activity | 3,50E-03 | CEWAF | GO_MolecularFunction-EBI-UniProt-GOA-ACAP-ARAP_16.06.2023_00h00 | opn1lw2, opn1mw1, opn1mw2, opn1sw1, opn1sw2, rho, rhol |
| photoreceptor activity | 3,97E-03 | LEWAF | GO_MolecularFunction-EBI-UniProt-GOA-ACAP-ARAP_16.06.2023_00h00 | opn1mw2, opn1sw1, rho |
